# Synthesis of versatile neuromodulatory molecules by a gut microbial glutamate decarboxylase

**DOI:** 10.1101/2024.03.02.583032

**Authors:** Pavani Dadi, Clint W. Pauling, Abhishek Shrivastava, Dhara D. Shah

## Abstract

Dysbiosis of the microbiome correlates with many neurological disorders, yet very little is known about the chemistry that controls the production of neuromodulatory molecules by gut microbes. Here, we found that an enzyme glutamate decarboxylase (*Bf*GAD) of a gut microbe *Bacteroides fragilis* forms multiple neuromodulatory molecules such as γ-aminobutyric acid (GABA), hypotaurine, taurine, homotaurine, and β-alanine. We evolved *Bf*GAD and doubled its taurine productivity. Additionally, we increased its specificity towards the substrate L-glutamate. Here, we provide a chemical strategy via which the *Bf*GAD activity could be fine-tuned. In future, this strategy could be used to modulate the production of neuromodulatory molecules by gut microbes.

## INTRODUCTION

Low levels of inhibitory neurotransmitter like γ-aminobutyric acid (GABA) have been associated with many different neurological disorders such as epilepsy, schizophrenia, autism spectrum disorder (ASD), ADHD, panic disorder, PTSD, major depressive disorder, progressive multiple sclerosis, dementia, and Alzheimer’s disease^1–12^. Besides GABA, taurine is another amino acid that is found in abundance in the brain^13^. Taurine is a GABA_A_ receptor agonist and exerts downstream effects similar to GABA^14^. Concentrations of taurine are altered in individuals with Alzheimer’s disease, where reduced levels of taurine are found in the brain^15–21^ whereas, higher urinary excretion of taurine is seen in elderly patients with dementia^22^. In a separate study, gnotobiotic mouse transplanted with fecal microbiota from an Alzheimer’s patient showed lower abundance of GABA and taurine in the feces^23^. Based on these observations, supplementation with taurine and its analogs has been investigated in Alzheimer’s disease models^24,25^. In one such study, Homotaurine, a molecule similar to taurine is shown to restore cognitive functions in patients with Alzheimer’s disease^26^. In addition, it was also observed that taurine reverses cognitive deficits in APP/PS1 mouse model^25^ and improves learning and memory in mice^24,27^.

Studies that correlate gut microbial dysbiosis to two of the prominent neurodegenerative disorders – Dementia and Alzheimer’s disease, often show modulations in the abundance of microbes of the genus *Bacteroides*^28–33^. *Bacteroides* are prominent members of the human gut^34,35^ and have the ability to produce GABA^36,37^. All *Bacteroides* encode the gene for the enzyme glutamate decarboxylase (GAD)^38^. Glutamate decarboxylases (GADs) are PLP (pyridoxal phosphate) dependent enzymes which catalyze conversion of an excitatory neurotransmitter glutamate to an inhibitory neurotransmitter GABA. However, very little is known about the chemistry of the glutamate decarboxylases encoded by *Bacteroides sp*.

Despite widespread prevalence of substrate promiscuity in glutamate decarboxylases (GADs) across different domains of life, substrate specificity and product formation landscape for most prokaryotic GADs, specifically GADs of *Bacteroides sp.* are still not clearly understood. For this reason, we explored the role of annotated GAD from *Bacteroides fragilis* in the production of neuromodulatory molecules like GABA, taurine and its analogs, and β-alanine. Our biochemical characterization of *Bf*GAD shows kinetic and functional divergence of this enzyme from other studied prokaryotic GADs. Our study shows that *Bf*GAD is not only specific to L-glutamate, but it can decarboxylate other substrates to produce multiple neuromodulatory molecules. Even within the substrate mix containing L-glutamate, we can detect products other than GABA. This points towards the ability of this enzyme to function in a complex system with multiple substrates which can be a beneficial trait for microbes present in the human gut. Through rational protein engineering, we have evolved *Bf*GAD which is capable of producing two-fold more taurine compared to the native *Bf*GAD. Additionally, we evolved *Bf*GAD to be more specific towards L-glutamate.

Based on our results of initial engineering with *Bf*GAD, the enzyme seems resilient in nature and is able to tolerate changes to the active site very well. We think that *Bf*GAD can be evolved with rational designing and engineering, either to produce various neuromodulatory molecules in specific ratios, or to synthesize a particular neuromodulatory molecule exclusively (Fig. 1a). This approach has the ability to generate variants of *Bf*GAD that might provide a road to therapeutic interventions for multiple neurodegenerative disorders.

**Fig. 1:**
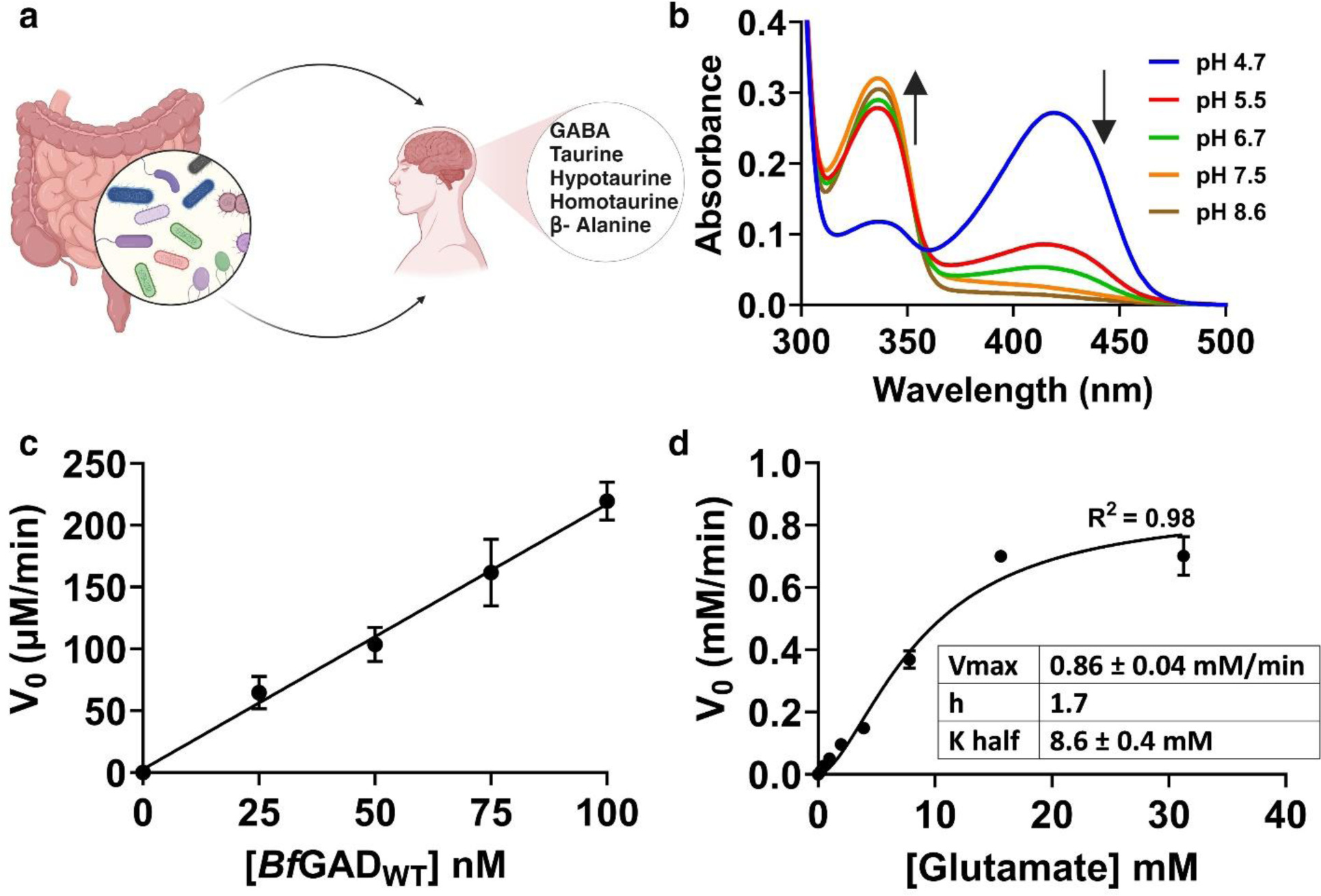
Characterization of *Bf*GAD_WT_. **a** Possible role of gut microbes in producing neuromodulatory molecules. **b** Spectra of PLP cofactor bound to the holoenzyme *Bf*GAD_WT_ captured at various pH, depicting tautomers of internal aldimine at 335 nm and 420 nm. **c** Plot of initial velocity as a function of *Bf*GAD_WT_ concentrations (n=4). **d** Plot of initial velocity vs. L-glutamate concentrations (n=2). The solid line is a sigmoidal fit to the data.

## RESULTS

### Glutamate Decarboxylases (GADs) are prevalent in the *Bacteroides* genus

A bioinformatics analysis showed that many species of the *Bacteroides* genus harbor genes annotated as GAD (Supplementary Fig. 1). We focused on a GAD (*Bf*GAD) from the model gut microbe, *Bacteroides fragilis*. Multiple sequence alignment performed with gut microbial GADs including *Bf*GAD revealed that multiple catalytic residues, either predicted to be involved in substrate binding (highlighted in cyan) or cofactor (PLP) binding (highlighted in yellow) are highly conserved (Supplementary Fig. 2)^39,40^. Moreover, in a phylogenetic analysis of *Bf*GAD with other gut microbial GADs, we see that the annotated glutamate decarboxylases from *Alistipes putredinis* and *Parabacteroides merdae* are closely related to *Bf*GAD (Supplementary Fig. 3).

### Wild-type *Bf*GAD (*Bf*GAD_WT_) forms an oligomer

The recombinant wild-type *Bf*GAD (*Bf*GAD_WT_) with an N-terminal His-tag was purified by Ni-NTA affinity chromatography. A protein band with an approximate expected mass of a *Bf*GAD_WT_ monomer (calculated mass 56 KDa) was observed with SDS PAGE, while a band indicating a tetrameric oligomer was observed with a native PAGE (Supplementary Fig. 4). Oligomeric composition of the *Bf*GAD_WT_, as determined by gel filtration chromatography, is presented in Supplementary Fig. 5a as chromatograms for the *Bf*GAD_WT_ at an acidic pH (blue) and a neutral pH (red). Peak 1 was in the void volume of the column indicating protein aggregates. There were two peaks representing two different oligomeric compositions (2 and 3) observed at each pH. Peak 2 was broad and corresponded to approximately 351 KDa, indicating a hexamer. Peak 3 was the major peak and exhibited an approximate molecular weight between a dimer and a tetramer at pH 4.7, whereas at pH 7.2 the peak 3 appeared to be closer to a dimer than a tetramer (Supplementary Fig. 5, a-c, 180 KDa at pH 4.7 and 150 KDa at pH 7.2) for *Bf*GAD_WT_. Our data indicated that *Bf*GAD_WT_ is an oligomeric enzyme which can either be a dimer or a tetramer. Furthermore, the addition of external PLP did not change the oligomeric state of the *Bf*GAD_WT_ (Supplementary Fig. 5d).

### PLP cofactor bound to the *Bf*GAD_WT_ undergoes pH dependent tautomeric changes

Spectral studies with *Bf*GAD_WT_ revealed that the enzyme was purified in its holo form with the covalently bound PLP cofactor. There were two absorption maxima observed for the *Bf*GAD_WT_ – 335 nm and 420 nm (Fig. 1b). These absorption maxima result from two tautomeric forms of the enzyme bound PLP cofactor, enolimine form at 335 nm and ketoenamine form at 420 nm^41–43^. The proportions of these tautomers vary depending on the pH (Fig. 1b). At lower pH, the species absorbing at 420 nm was more dominant, whereas at higher pH, 340 nm absorbing form became more prevalent. With increase in pH from 4.7 to 8.6, there was a decrease in 420 nm absorbing species with concomitant increase in 340 nm absorbing species (Fig. 1b). These changes in the absorbances of the bound PLP cofactor as the function of the change in pH with *Bf*GAD_WT_, have been seen with other microbial GADs^44–46^. The analysis of the pH versus absorbance curve for the 420 nm wavelength indicated that the transition in absorbance in response to pH variation is influenced by the involvement of multiple protons (Supplementary Fig. 6a). Previously with *E. coli* GAD, Tramonti et al. and Capitani et al. showed that the less hydrated active site at neutral pH drove the formation of enolimine tautomer (340 nm) whereas more polar active site at acidic pH predominantly produced ketoenamine tautomer (420 nm)^39,47^. Additionally, the alterations in cofactor absorbance corresponding to different pH were in close agreement with the *Bf*GAD_WT_’s activity profiles measured across the same range of pH values. Particularly, at pH 4.7, where *Bf*GAD_WT_ with a bound PLP predominantly absorbed at 420 nm, the enzyme also exhibited its maximum catalytic activity (Supplementary Fig. 6b). Additionally, enzymatic activity was found to be minimal at a pH other than 4.7 (Supplementary Fig. 6b). Therefore, all subsequent activity assays were carried out at pH 4.7.

### Evidence for allosteric regulation of *Bf*GAD_WT_

The activity assays conducted with varying *Bf*GAD_WT_ concentrations exhibited a linear relationship between the initial reaction velocities and enzyme concentrations (Fig. 1c). From the plot of enzyme concentration variation, we found an optimal *Bf*GAD_WT_ concentration for subsequent kinetic assays. Interestingly, while varying L-glutamate concentrations, the plot of substrate concentration versus initial velocity displayed a sigmoidal curve instead of a typical hyperbolic dependence, suggestive of an allostery in *Bf*GAD_WT_. This is an example of a homotropic allosteric regulation. From the fit of this plot with the hill equation (eq. 2), the analyzed K_0.5_ (K_half_) was 8.6 ± 0.4 mM and V_max_ was 0.86 ± 0.04 mM/min with a hill coefficient of 1.7 (Fig. 1d). (K_half_)^h^ is also known as K_prime_ (K’), which is equivalent to K_m_ from a hyperbolic kinetics of non-allosteric enzymes, when h=1. However, in cooperative systems where h ≠1 like presented here for *Bf*GAD_WT_ then, K’ (K_half_^h^) no longer represents the substrate concentration required to achieve half of maximal velocity.

### Engineered *Bf*GAD variants exhibit structural and catalytic perturbations

The earliest microbial GAD structures were solved for *E. coli* GADs (GAD*A*, PDB ID 1XEY and GAD*B*, PDB ID 1PMM) (Supplementary Fig. 7)^39,40^. Both structures were solved with the bound ligand in the active site. Based on the interactions of the acetate ion in the GAD*B* active site and glutarate (substrate analog) in the GAD*A* active site (Supplementary Fig. 7b), we selected residues for the evolution of the *Bf*GAD_WT_ via rational design^39,40,48^. Two such residues that are present in the active sites of both *Ec*GADs are phenylalanine 63 (corresponds to F81 in *Bf*GAD_WT_) and aspartate 86 (corresponds to D104 in *Bf*GAD_WT_) that make H-bonding interactions with one of the carboxylates of the substrate analog and an acetate ion^39,40^. Specifically, in *E. coli* GAD*B* ligand bound structure, the carboxylate group of acetate received a hydrogen bond from the amide nitrogen of the F63 (F81 *Bf*GAD_WT_) and a side chain carboxylate of D86 (D104 *Bf*GAD_WT_) of the neighboring subunit^39^. Whereas in *E. coli* GAD*A* ligand bound structure, one of the carboxylate group of the glutarate (substrate analogue) forms H-bond with the amide nitrogen of F63 (F81 *Bf*GAD_WT_) and with the side chain carboxylate of D86 (D104 *Bf*GAD_WT_) of the neighboring subunit^40^. These residues are completely conserved in annotated GADs from prominent gut microbes with the exception of *Eggerthella lenta* that harbors shorter GAD with the absence of a conserved phenylalanine (Supplementary Fig. 2). Most microbial GADs are functionally active as dimers, with residues from both monomers contributing to the active site^39,40,44,49^. Recently, structure of glutamate decarboxylase from *Bacteroides thetaiotamicron* (*Bt*GAD), bound with a substrate analog (glutarate) was solved^44^. With these available structures, we aligned the substrate analog (glutarate) bound structures of *Ec*GAD*A* (PDB ID 1XEY) and *Bt*GAD (PDB ID 7X51) with *Bf*GAD_WT_ dimer created with AlphaFold2 (Fig. 2). Based on these structural alignments we engineered two enzymes by introducing single amino acid alterations at two separate positions to test if the substrate preference of the *Bf*GAD_WT_ can be evolved. These changes were Asp104Asn (D104N) and Phe81Trp (F81W). Figure 2a and 2b illustrate the locations of these residues within the active site of *Bf*GAD_WT_ (green). These residues are depicted in close proximity to glutarate and the cofactor PLP bound in the active site of *Ec*GAD*A*, which has been superimposed on the *Bf*GAD_WT_ dimer (Fig. 2a, 2b). Similarly, Fig. 2c and 2d display aligned *Bf*GAD_WT_ and *Bt*GAD structures with glutarate and PLP bound in the active site of *Bt*GAD. In addition to *Bf*GAD_WT_, the structures of variants, *Bf*GAD_D104N_ (Fig. 2a, 2c) and *Bf*GAD_F81W_ (Fig. 2b, 2d) were then superimposed with ligand bound structures of *Ec*GAD and *Bt*GAD.

**Fig. 2:**
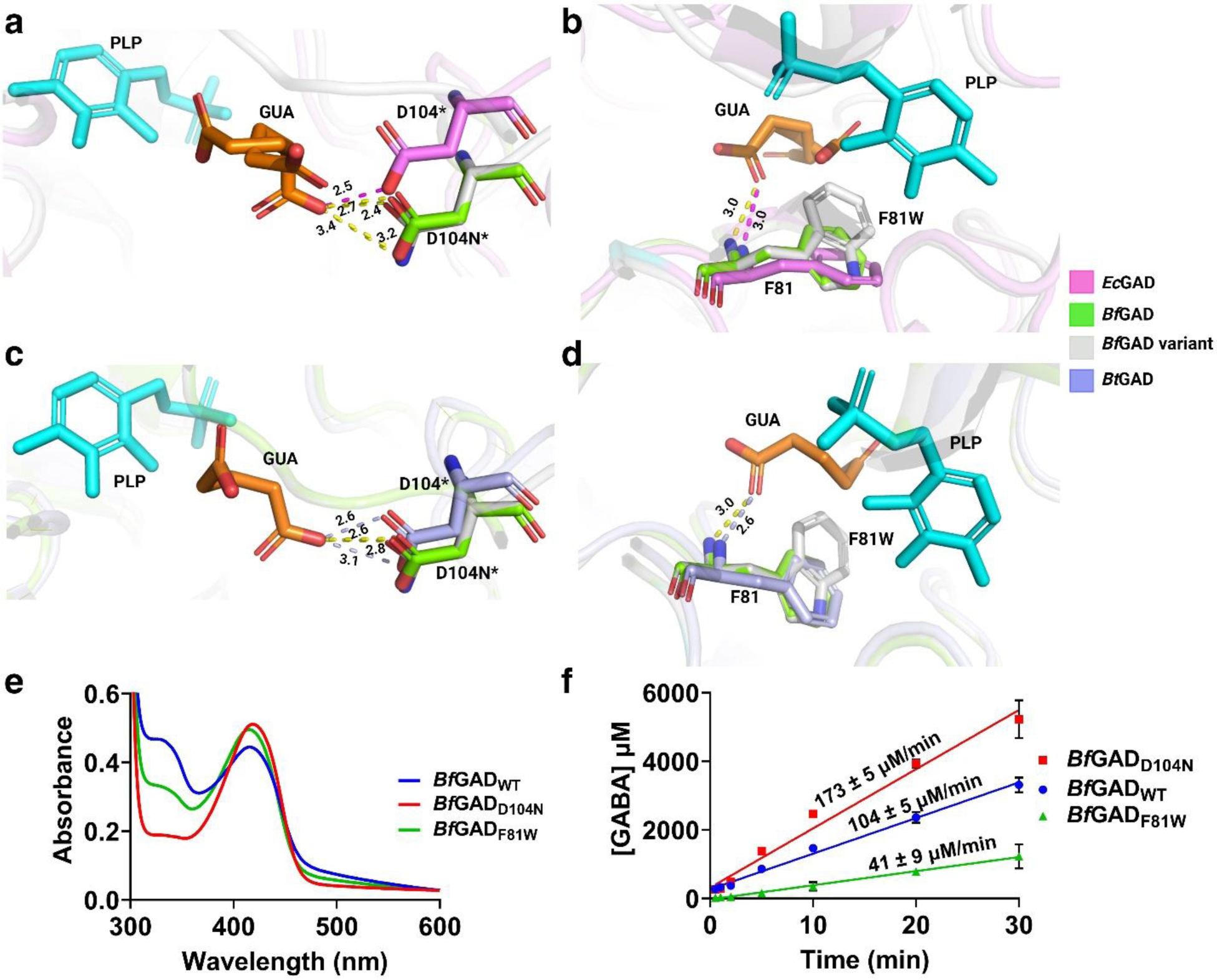
Structural and catalytic perturbations of engineered *Bf*GADs. **a** Aligned structures of *Ec*GAD (1XEY, pink), *Bf*GAD_WT_ (model, green) illustrating the position of the residue D104* (*residue from the neighbouring monomer), and *Bf*GAD_D104N_ depicting the point substitution D104N*(model, gray). **b** Aligned structures of *Ec*GAD (1XEY, pink), *Bf*GAD_WT_ (model, green) illustrating the position of the residue F81, and *Bf*GAD_F81W_ depicting the point substitution F81W (model, gray). **c** Aligned structures of *Bt*GAD (7X51, purple), *Bf*GAD_WT_ (model, green) illustrating the position of the residue D104* (*residue from the neighbouring monomer), and *Bf*GAD_D104N_ depicting the point substitution D104N*(model, gray). **d** Aligned structures of *Bt*GAD (7X51, purple), *Bf*GAD_WT_ (model, green) illustrating the position of the residue F81, and *Bf*GAD_F81W_ depicting the point substitution F81W (model, gray). The dashed lines indicate hydrogen bond interactions of the amino acid residues with substrate analog, glutarate (GUA) where pink bonds are for interactions between *Ec*GAD-GUA, yellow bonds are for interactions between *Bf*GAD-GUA, and purple bonds are for interactions between *Bt*GAD-GUA. **e** Spectra of PLP cofactor bound to the holoenzymes *Bf*GAD_WT_ and engineered variants, depicting tautomers of internal aldimine at 335 nm and 420 nm. **f** Progress curves depicting initial velocities for reactions catalyzed *Bf*GAD_WT_ and variants (n=3).

The overall active site architecture of *Bf*GAD is mostly consistent with that of *Ec*GAD and *Bt*GAD; however, notable positional shifts were identified among the residues of interest. Specially, aspartic acid (D104) showed a considerable difference in *Ec*GAD versus *Bf*GAD active site. Additionally, we detected changes in the H-bonding interactions between the substrate analog and residues of interest when substitutions were made to create engineered *Bf*GAD variants. We were able to purify both variants, *Bf*GAD_D104N_ and *Bf*GAD_F81W_ with the same conditions as the *Bf*GAD_WT_. Both of these variants were active and purified as holoenzymes (PLP bound form). Additionally, we collected UV-Vis spectra of *Bf*GAD_WT_ and both variants at pH 4.7 (Fig. 2e). While each of the three enzyme preparations exhibits absorbance maxima at both 335 nm and 420 nm, the relative amounts of these two tautomeric forms differ across the enzymes. Furthermore, we measured initial velocities in the presence of the native substrate L-glutamate via progress curve analysis for engineered *Bf*GADs along with *Bf*GAD_WT_. With that, we detected variations in the decarboxylation reaction rates across these enzymes (Fig. 2f). *Bf*GAD_D104N_ showed 1.7-fold increase whereas *Bf*GAD_F81W_ showed 2.5-fold decrease in the initial velocities compared to the *Bf*GAD_WT_ for decarboxylation reactions with L-glutamate. These observed differences in catalysis between the wild-type and engineered variants of the *Bf*GAD due to the structural perturbations resulting from amino acid substitutions at the active site, reinforce the role of F81 and D104 in establishing interactions with the substrate.

### Wild-type and engineered *Bf*GADs can decarboxylate substrates other than L-glutamate

As mentioned above, the purified *Bf*GAD_WT_ is able to catalyze the conversion of L-glutamate to GABA. To test if it can utilize the D-form of the glutamate as a substrate, we conducted activity assays with D-glutamate using the GABase assay system and TLC. Our results show that both the wild-type and engineered *Bf*GADs were unable to decarboxylate D-glutamate (Supplementary Fig. 8). Most microbial GADs studied so far show high substrate specificity towards L-glutamate^49–52^. However, there are examples of archaeal GADs that prefer L-aspartate over L-glutamate^53,54^. An archaeal GAD from *P. horikoshii* also shows decarboxylation activity with L-cysteate^53^. In addition to archaeal GADs, certain mammalian glutamate decarboxylase (GAD) homologs are known to decarboxylate one or more of the non-native substrates such as L-aspartate, L-cysteate, and L-cysteine sulfinate^55,56^. Moreover, proteins similar to GADs have been demonstrated to form taurine in some marine microbes^57^. However, there are limited systematic studies investigating the production of taurine, its analogs, and β-alanine using L-CA (cysteate), L-CSA (cysteine sulfinate), L-HCA (homocysteate), and L-aspartate as substrates (Scheme 1), specifically for gut microbial GADs. Based on two key observations, we hypothesized that gut microbial GADs might utilize these molecules as substrates: 1. *E. coli* GAD can accept the phosphonated form of L-glutamate as a substrate^58^. and 2. Archaeal and eukaryotic GADs exhibit diverse substrate specificities, enabling them to catalyze the decarboxylation reactions of non-native substrates^53,54,57,59,60^.

If *Bf*GAD_WT_ or engineered *Bf*GADs are able to decarboxylate substrates other than the native substrate L-glutamate, then the detection of the common product CO_2_ will be a positive test for the utilization of other substrates (Scheme 1). For this reason, we decided to use a headspace GC to measure the CO_2_ evolved from reactions catalyzed by either *Bf*GAD_WT_ or engineered *Bf*GADs during the decarboxylation of various substrates. Fig. 3 shows data collected for *Bf*GAD_WT_ and engineered *Bf*GADs with five different substrates via headspace GC. Panels 3a, 3c, and 3e demonstrate chromatograms with the CO_2_ peaks, visible immediately after 12.8 min, produced by the reactions of *Bf*GAD_WT_, *Bf*GAD_D104N_, and *Bf*GAD_F81W_ respectively with five different substrates. Whereas 3b, 3d, and 3f depict CO_2_ peak areas from the decarboxylation reactions of the substrates after 24 h incubation with either *Bf*GAD_WT_ or engineered *Bf*GADs. Here, *Bf*GAD_WT_ showed significant production of CO_2_ generated from the decarboxylation of both L-glutamate (blue) and L-CSA (red) (Fig. 3a, 3b). This is the first indication to our knowledge of a gut microbial GAD, specifically a GAD from *Bacteroides,* that can utilize L-CSA as a substrate. In addition to L-CSA, *Bf*GAD_WT_ was capable of decarboxylating L-HCA (orange) and L-CA (green) to small extents as evidenced by the small CO_2_ peaks (Fig. 3a) in the chromatogram and small peak areas for evolved CO_2_ (Fig. 3b). Additionally, *Bf*GAD_WT_ was able to decarboxylate L-aspartate (purple) (Fig. 3a, 3b). For *Bf*GAD_WT_, if CO_2_ production peak area with the native substrate L-glutamate is considered to be 100% then we observed 54%, 3%, 6% and 20% of CO_2_ production with L-CSA, L-CA, L-HCA, and L-Asp respectively within 24 h which increased to 75% (L-CSA), 6% (L-CA), 7% (L-HCA), and 28% (L-Asp) within 48 h compared to the native substrate L-glutamate (Supplementary Fig. 9a).

**Fig. 3:**
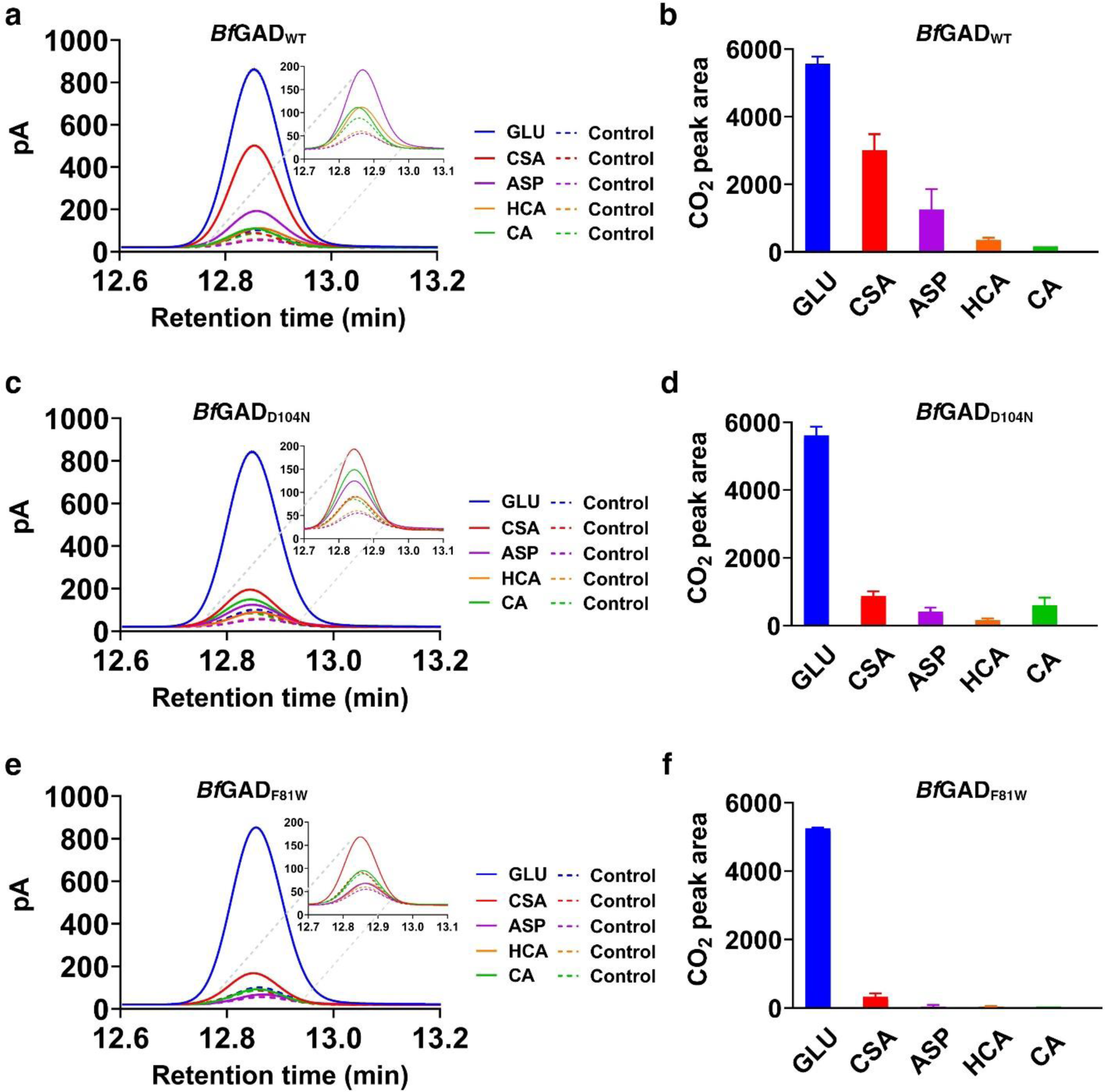
Wild-type and engineered *Bf*GADs can decarboxylate multiple substrates. Chromatograms showing the CO_2_ peaks obtained from the decarboxylation reactions of multiple substrates after 24 h incubation with **a** *Bf*GAD_WT_, **c** *Bf*GAD_D104N,_ and **e** *Bf*GAD_F81W_ using headspace GC (n=3, only one representative CO_2_ peak is depicted for every reaction). The CO_2_ peak areas are depicted for the decarboxylation of various substrates after 24 h incubation with **b** BfGAD_WT_, **d** BfGAD_D104N,_ and **f** BfGAD_F81W_ (n=3). Various substrates are presented as specific colors which are consistent in all panels where L-glutamate is blue, L-CSA is red, L-Asp is purple, L-HCA is orange and L-CA is green.

For *Bf*GAD_D104N_, CO_2_ production is comparable to the *Bf*GAD_WT_ while using native substrate L-glutamate. Compared to the peak area of CO_2_ production for L-glutamate (native substrate), we observed around 16%, 11%, 3% and 7% CO_2_ production from L-CSA, L-CA, L-HCA, and L-aspartate respectively within 24 h (Fig. 3c, 3d) that increased to 18% for both L-CSA and L-CA, 7% for L-HCA and slightly decreased with L-aspartate within 48 h (Supplementary Fig. 9b). Based on the data collected with *Bf*GAD_WT_ and *Bf*GAD_D104N_, residue Asp104 (D104) from the neighboring monomer is playing a crucial role in accommodating various substrates. This engineered enzyme was also able to decarboxylate L-CSA, L-HCA, and L-Asp in addition to the native substrate L-glutamate but less efficiently than the *Bf*GAD_WT_, specifically the activity towards L-CSA was significantly impacted. It is likely that the negative charge of the side chain carboxylate from D104 is important for the specificity towards alternative substrates L-CSA, L-HCA and L-Asp but not for the native substrate L-glutamate. As a result, when the charge was removed due to the substitution of Asp (D) to Asn (N) at position 104, the catalysis with these alternate substrates was affected, but not with the native substrate L-glutamate. Interestingly, we noticed a 2-2.5-fold increase in the production of taurine with this evolved enzyme compared to the *Bf*GAD_WT_ (Fig. 3d), indicating that the substitution from D to N was favorable for taurine production.

The second engineered enzyme, *Bf*GAD_F81W_ retained its activity with the native substrate L-glutamate as well but as observed in the initial velocity experiments (Fig. 2f), it was slower compared to *Bf*GAD_WT_. If the CO_2_ peak area of *Bf*GAD_WT_ or *Bf*GAD_D104N_ with the native substrate L-glutamate is considered to be 100%, *Bf*GAD_F81W_ showed the peak area of around 93-94% with the same concentration of the substrate. Additionally, there was minimal decarboxylation activity of *Bf*GAD_F81W_ with L-CA, L-HCA, and L-ASP. However, this evolved enzyme still retained considerable decarboxylation activity with L-CSA. It is interesting to note that a bulky substitution of tryptophan (W) at position 81, instead of phenylalanine (F) (Fig. 2b, 2d), made the enzyme more specific towards the native substrate L-glutamate. Based on the CO_2_ evolution captured by our GC data, the substrate preference for *Bf*GAD_WT_ was L-Glu > L-CSA > L-Asp > L-HCA > L-CA that changed to L-Glu > L-CSA ≥ L-CA > L-Asp > L-HCA for *Bf*GAD_D104N_. Whereas for *Bf*GAD_F81W_, only L-glutamate and L-CSA showed decarboxylation and L-glutamate was a much better substrate than L-CSA.

### Wild-type and evolved *Bf*GADs produce multiple neuromodulatory molecules

The generation of CO_2_ in reactions facilitated by *Bf*GAD_WT_ and its engineered variants prompted us to detect and identify products resulting from decarboxylation with various substrates. Both *Bf*GAD_WT_ and *Bf*GAD_D104N_ produced decarboxylated products – hypotaurine, taurine, homotaurine, and β-alanine from substrates L-CSA, L-CA, L-HCA, and L-Asp respectively (Fig. 4, a-d, Supplementary Fig. 10, a-d). Intensity plots generated from TLC plates showed that *Bf*GAD_WT_ decarboxylated L-CSA and L-Asp to produce hypotaurine and β-alanine more efficiently than *Bf*GAD_D104N_ (Fig. 4a, 4c). However, *Bf*GAD_D104N_ was better at decarboxylating L-CA to taurine than *Bf*GAD_WT_ (Fig. 4b). We do not observe much of decarboxylated product taurine with *Bf*GAD_WT_ from L-CA decarboxylation during a 24 h incubation, whereas *Bf*GAD_D104N_ shows a significant production of taurine during this time frame (Fig. 4b, Supplementary Fig. 10e). While our CO_2_ evolution experiments have shown that *Bf*GAD_WT_ can catalyze L-CA decarboxylation in 24 h, the failure to observe decarboxylated product on TLC may be attributed to the low concentration of taurine produced, making it undetectable on a TLC plate. Both enzymes *Bf*GAD_WT_ and *Bf*GAD_D104N_ were able to catalyze the decarboxylation of L-HCA to homotaurine to a very small extent (Fig. 4d). We also verified that the product from the L-Asp decarboxylation was β-alanine and not L-alanine. Although, the R*_f_* (retention factor) values were almost similar for β-alanine and L-alanine, the staining with the ninhydrin differs for these molecules. β-alanine exhibited a purple color whereas L-alanine shows a brick red color with ninhydrin stain (Supplementary Fig. 10c). We did not find any activity of *Bf*GAD_F81W_ with L-CA, L-Asp, and L-HCA and were unable to detect taurine, β-alanine, and homotaurine even after 48 h incubation (Supplementary Fig. 11a, 11b). These results reinforce our observations from CO_2_ evolution experiments. Additionally, despite the detection of CO_2_ during the decarboxylation activity of *Bf*GAD_F81W_ with L-CSA (Fig. 3f), we could not observe detectable hypotaurine spots on the TLC plate (Supplementary Fig. 11a). We hypothesize that the amount of hypotaurine produced by *Bf*GAD_F81W_ may not be sufficient to be detected via TLC, as a higher concentration of molecules (in mM range) are needed for TLC analysis.

**Fig. 4:**
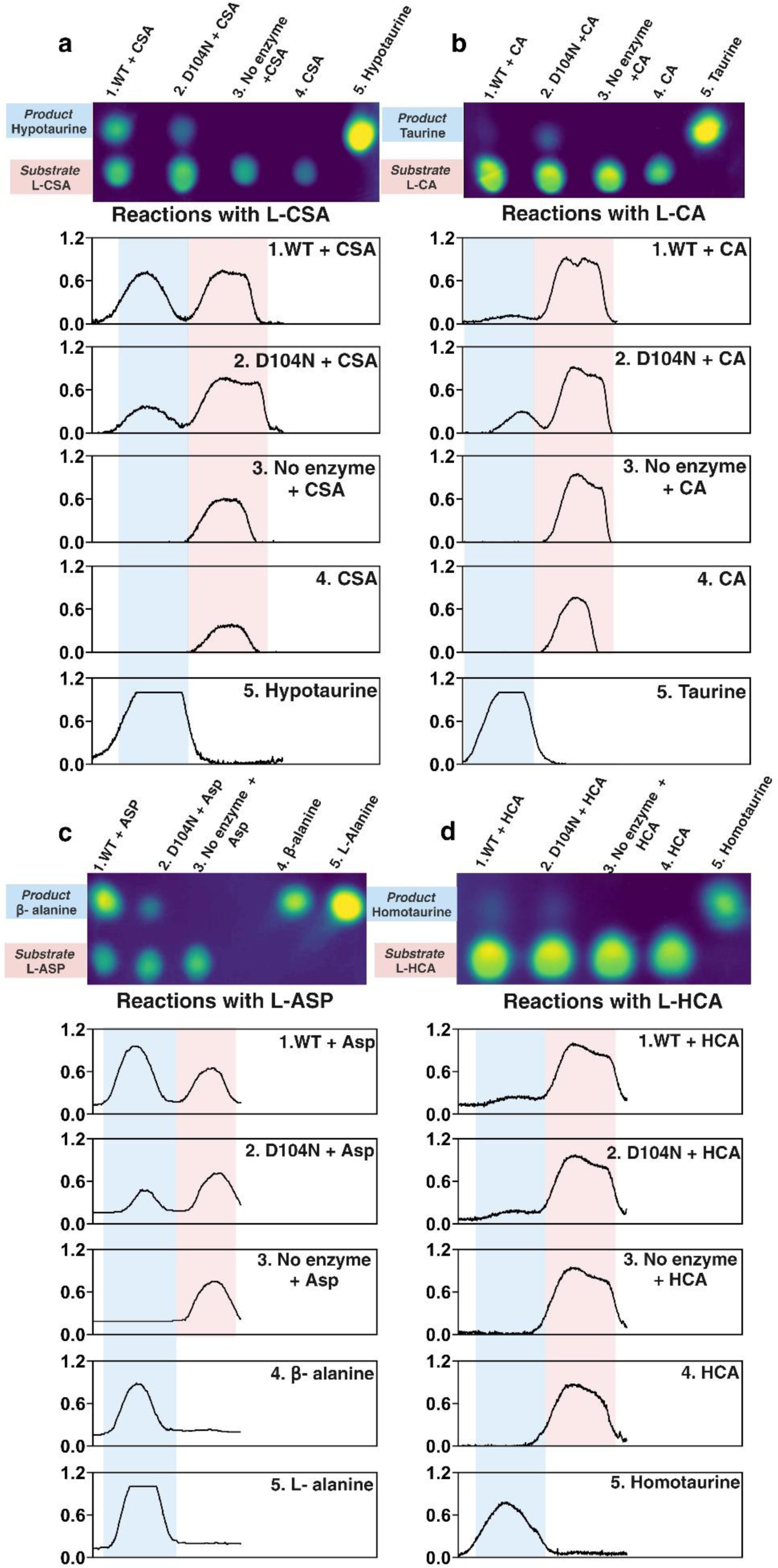
Production of multiple neuromodulatory molecules by Wild-type and evolved *Bf*GADs. Intensity analysis of TLC plates depicting intensity plots for each lane. The peak size is proportional to the intensity of the sample spot on the TLC plate. **a** Production of hypotaurine from the decarboxylation of L-CSA catalyzed by *Bf*GAD_WT_ and *Bf*GAD_D104N_ in 24 h as indicated by TLC intensity analysis. For intensity plots, peaks highlighted in pink are for the substrate L-CSA and peaks highlighted in blue are for the product hypotaurine. **b** Production of taurine from the decarboxylation of L-CA catalyzed by *Bf*GAD_WT_ and *Bf*GAD_D104N_ in 24 h as indicated by TLC intensity analysis. For intensity plots, peaks highlighted in pink are for the substrate L-CA and peaks highlighted in blue are for the product taurine. **c** Production of β-alanine from the decarboxylation of L-ASP catalyzed by *Bf*GAD_WT_ and *Bf*GAD_D104N_ in 24 h as indicated by TLC intensity analysis. For intensity plots, peaks highlighted in pink are for the substrate L-ASP and peaks highlighted in blue are for the product β-alanine. **d** Production of homotaurine from the decarboxylation of L-HCA catalyzed by *Bf*GAD_WT_ and *Bf*GAD_D104N_ in 24 h as indicated by TLC intensity analysis. For intensity plots, peaks highlighted in pink are for the substrate L-HCA and peaks highlighted in blue are for the product homotaurine. The original TLC plates are shown in Supplementary Fig. 10.

### *Bf*GAD_WT_ and *Bf*GAD_D104N_ generate neuromodulatory molecules at differential abundances, even when presented with a mixed substrate pool

In a complex gut environment, the organism will encounter multiple substrates simultaneously. To investigate how *Bf*GAD_WT_ functions in such a complex environment, we conducted competition assays between various substrates. To this end, we tried to detect the presence of decarboxylated products in mixtures containing the native substrate L-glutamate with a specific alternate substrate. Our data with *Bf*GAD_WT_ indicate that when the concentration ratio of the native substrate to the alternative substrate was 1:1, GABA was predominantly seen as the primary decarboxylated product (Supplementary Fig. 12f). However, when the concentration ratios were changed to 1:5 (L-Glu with L-CSA/L-Asp) and 1:10 (L-Glu with L-CSA/L-CA/L-Asp), a gradual increase over time in the formation of the alternative decarboxylated products - hypotaurine, taurine, and β-alanine was observed (Fig. 5-7). For the *Bf*GAD_WT_ catalyzed decarboxylation reactions of L-CSA and L-Asp, the resulting products hypotaurine and β-alanine could be observed when these substrates were in 5-fold excess to the native substrate L-glutamate. However, the accumulation of products hypotaurine and β-alanine becomes significant only after 9 hours (for β-alanine) – 24 h (for hypotaurine) (Supplementary Fig. 12d, 12e). In contrast, when L-CSA and L-Asp are used in 10-fold excess of L-glutamate, the resulting products, hypotaurine and β-alanine (peaks highlighted in pink) started accumulating significantly much earlier around 3 hours (Fig. 5, 7). We did not observe the accumulation of taurine even when 10-fold excess of L-CA was mixed with L-glutamate in the *Bf*GAD_WT_ catalyzed reaction (Fig. 6). However, as mentioned earlier, we were able to detect CO_2_ in the same time frame (48 h) in the reaction catalyzed by *Bf*GAD_WT_ when L-CA was provided as the sole substrate (Supplementary Fig. 9a). The inability to detect taurine might be because of the low concentrations produced under the competitive environment of the mixed substrate pool. Unlike *Bf*GAD_WT,_ *Bf*GAD_D104N_ was able to decarboxylate and accumulate taurine (peaks highlighted in pink) to a small extent when L-CA is mixed with L-glutamate at a 10-fold excess concentration, specially between 24-48 h (Fig. 6). Interestingly, in all mixed substrate experiments, we observed maximum accumulation of the native product GABA (peaks highlighted in blue) within 3 h. The intensity peaks showed no further increase after 3 h time point. This shows that even in the presence of other substrates the native activity of L-glutamate decarboxylation occurred at the highest velocity for all *Bf*GADs compared to the decarboxylation reactions of other substrates.

**Fig. 5:**
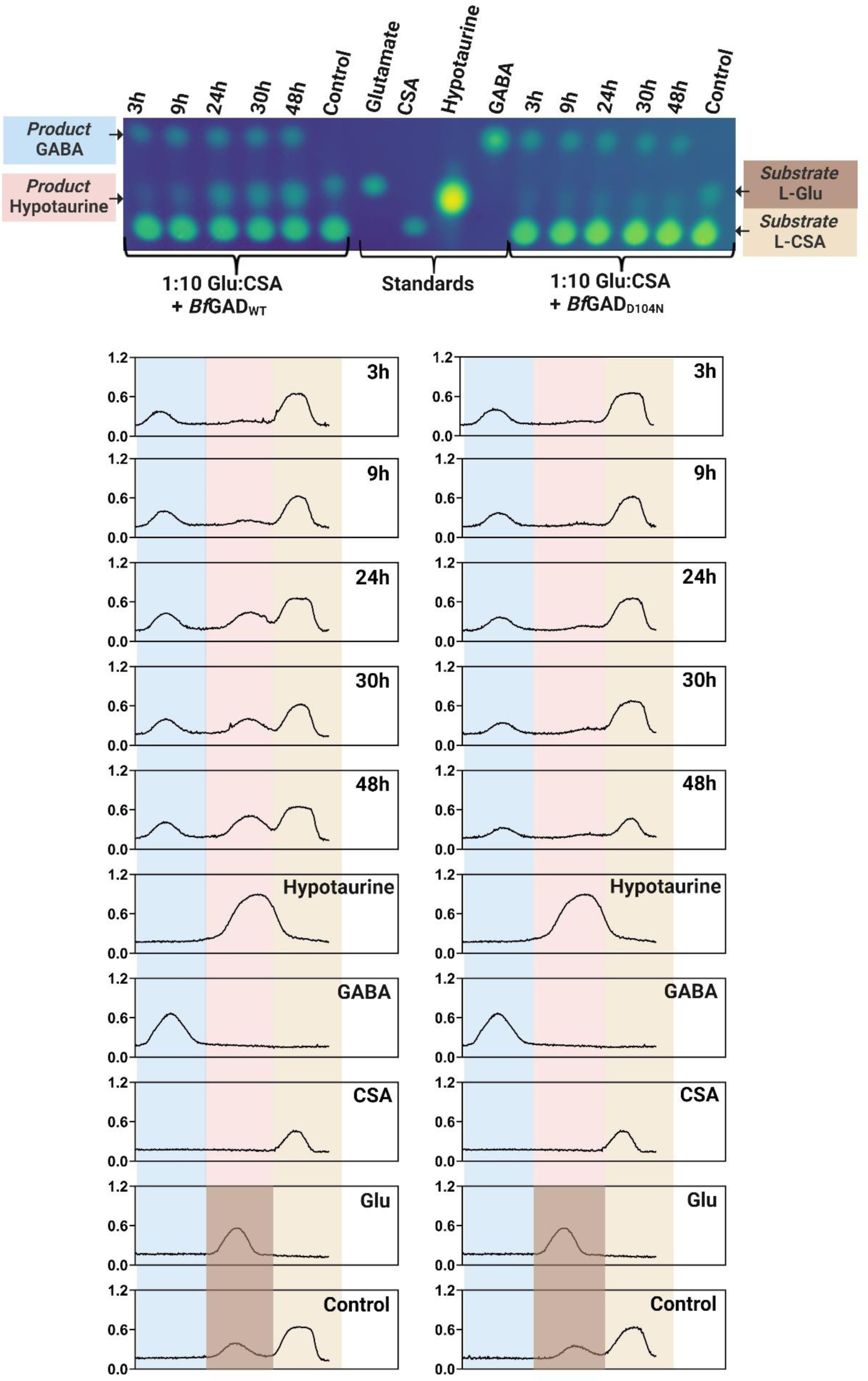
*Bf*GAD_WT_ and *Bf*GAD_D104N_ generate hypotaurine when presented with a mixture of L-glutamate and L-CSA. Intensity analysis of TLC plates for various timepoints of decarboxylation reactions catalyzed by *Bf*GAD_WT_ and *Bf*GAD_D104N_ for mixed substrates where the concentration ratio of L-glutamate to L-CSA is 1:10. Standards of L-glutamate, L-CSA, GABA, and hypotaurine were included as references. Production of hypotaurine (peaks highlighted in pink) can be seen as the peak area increases over time. GABA peaks are highlighted in blue, peaks for the substrate L-CSA are highlighted in beige, and peaks for the substrate L-glutamate are in brown. Spots visible on TLC plates from top to bottom show corresponding intensity peaks from left to right.

**Fig. 6:**
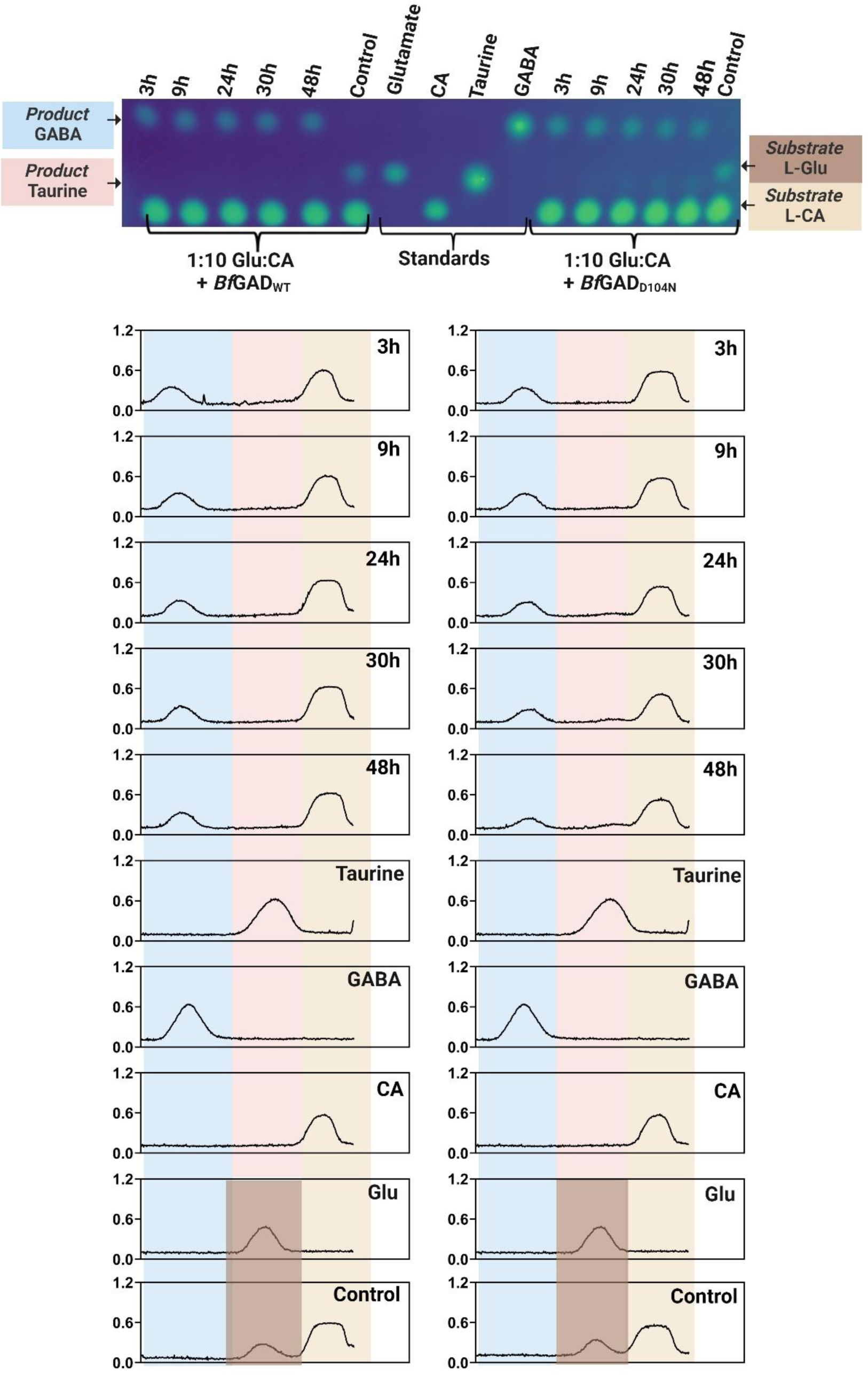
*Bf*GAD_D104N_ generate taurine when presented with a mixture of L-glutamate and L-CA. Intensity analysis of TLC plates for various timepoints of decarboxylation reactions catalyzed by *Bf*GAD_WT_ and *Bf*GAD_D104N_ for mixed substrates where the concentration ratio of L-glutamate to L-CA is 1:10. Standards of L-glutamate, L-CA, GABA, and taurine were included as references. Production of taurine (peaks highlighted in pink) can be seen as the peak area increases over time for *Bf*GAD_D104N_ catalyzed reactions. GABA peaks are highlighted in blue, peaks for the substrate L-CA are highlighted in beige, and peaks for the substrate L-glutamate are in brown. Spots visible on TLC plates from top to bottom show corresponding intensity peaks from left to right.

**Fig. 7:**
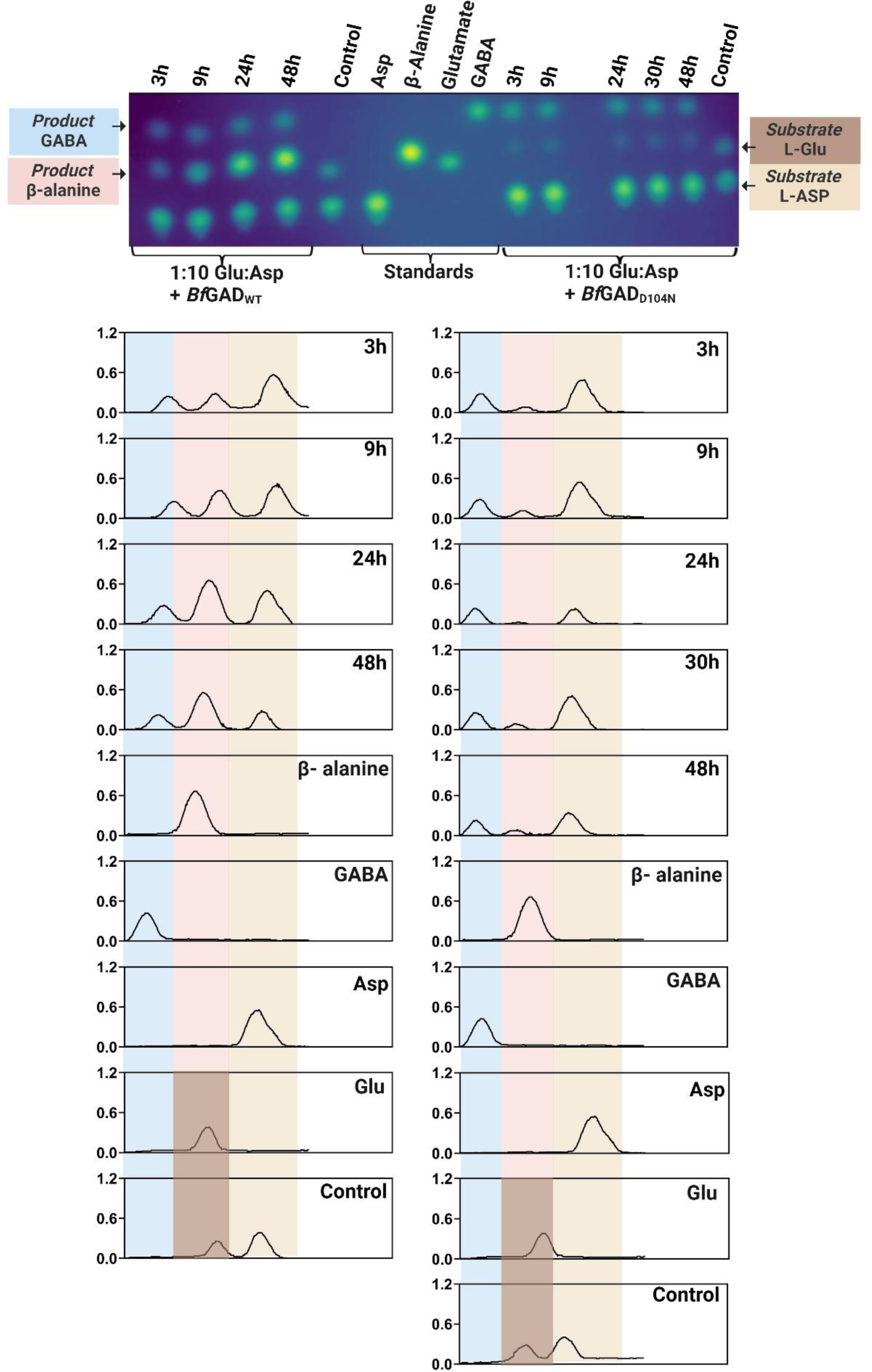
*Bf*GAD_WT_ and *Bf*GAD_D104N_ generate β-alanine when presented with a mixture of L-glutamate and L-aspartate. Intensity analysis of TLC plates for various timepoints of decarboxylation reactions catalyzed by *Bf*GAD_WT_ and *Bf*GAD_D104N_ for mixed substrates where the concentration ratio of L-glutamate to L-aspartate is 1:10. Standards of L-glutamate, L-aspartate, GABA, and β-alanine were included as references. Production of β-alanine (peaks highlighted in pink) can be seen as the peak area increases over time. The increase over time for *Bf*GAD_D104N_ is very clear from 3 h to 9 h. Due to the differences in the background intensity of the TLC plate in areas containing 3 h and 9 h samples vs areas containing 24 h, 30 h and 48 h samples from *Bf*GAD_D104N_ catalyzed reaction, the increase is the intensity peak is not uniform. GABA peaks are highlighted in blue, peaks for the substrate L-aspartate (L-ASP) are highlighted in beige, and peaks for the substrate L-glutamate are in brown. Spots visible on TLC plates from top to bottom show corresponding intensity peaks from left to right.

Additionally, the decarboxylated products hypotaurine and taurine were analyzed using LC-ESI-MS/MS in addition to TLC to confirm their presence and mass in mixed substrate assays where L-glutamate to L-CSA or L-CA ratios were 1:10. The qualitative analysis of these molecules was conducted using MRM mode by running mixed standards (Supplementary Fig. 13). Figure 8a depicts the presence of the product hypotaurine and the remaining L-CSA substrate in the mixed substrate reactions with a 1:10 ratio of L-glutamate:L-CSA catalyzed by *Bf*GAD_WT_. Figure 8b depicts the presence of taurine and remaining L-CA in the mixed substrate reactions with a 1:10 ratio of L-Glutamate:L-CA catalyzed by *Bf*GAD_D104N_. We did not analyze the native substrate and product, L-glutamate and GABA in these samples using LC-MS/MS because the presence of these molecules was confirmed by both TLC and GABase assay prior to LC-MS/MS analysis.

**Figure 8:**
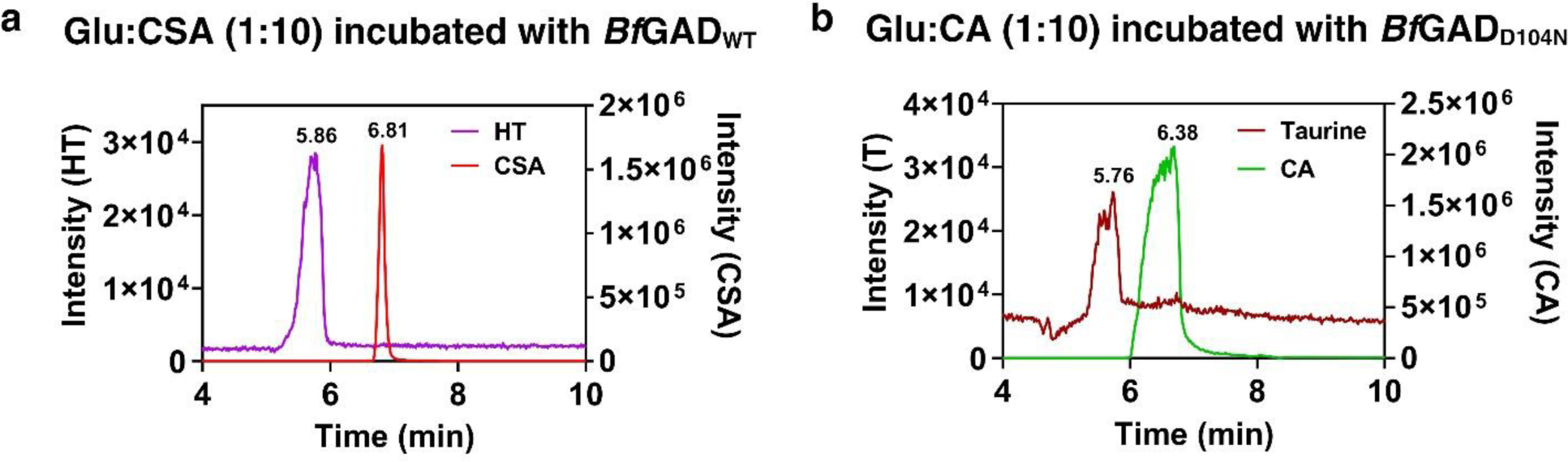
Hypotaurine and Taurine detection via LC-MS/MS from *Bf*GAD_WT_ and *Bf*GAD_D104N_ catalyzed mixed substrate reactions. **a** Ion chromatograms of product hypotaurine and the remaining L-CSA substrate in the mixed substrate reactions with a 1:10 ratio of L-glutamate:L-CSA catalyzed by *Bf*GAD_WT_. **b** Ion chromatograms of taurine and remaining L-CA in the mixed substrate reactions with a 1:10 ratio of L-Glutamate:L-CA catalyzed by *Bf*GAD_D104N._

## DISCUSSION

Prior research has implicated gut microbial contributions to the production of γ-aminobutyric acid (GABA), specifically from *Bacteroides* genus^36,37^. GABA, a neurotransmitter, is modulated in many neurodegenerative diseases, including Alzheimer’s and dementia, where lower GABA levels have been consistently reported^12^. Microbes from the genus *Bacteroides* are known to be fluctuated in individuals afflicted by Alzheimer’s and dementia^30–33^. Due to the prevalent nature of *Bacteroides* in the gut and the modulation of Bacteroides during Alzheimer’s, we decided to understand the mechanism by which GABA is produced in these organisms. For this reason, an annotated glutamate decarboxylase from *B. fragilis* (*Bf*GAD) was selected as a candidate enzyme representing annotated GADs in all *Bacteroides*.

Most of the studied microbial GADs are oligomeric which include dimeric, tetrameric or hexameric states^39,44,49,61–63^. It is known that the functional GAD unit is a dimer as residues from both monomers make an intact active site and play an important role during the catalysis. Therefore, an oligomeric composition that is not an even number would be unusual in this context. Our gel filtration chromatography data with pH 4.7 buffer gave a molecular weight of the protein that is higher than a dimer but lower than a tetramer. We hypothesize that the protein likely exists in a dynamic equilibrium between these two states, with a tendency towards a tetrameric state. We observed an intermediate molecular weight possibly due to the inability to resolve dimeric and tetrameric states completely. However, at pH 7.2, there might be a shift in this equilibrium favoring a more dimeric state which is observed with a molecular weight closer to that of a dimer than a tetramer. This shift in oligomeric states has been observed previously in *E. coli* GAD that undergoes significant changes in oligomeric forms upon shift in the pH. Thus, purified *Bf*GAD_WT_ behaves similarly in the oligomeric composition to other characterized prokaryotic GADs. Our data suggests that there might be a dynamic equilibrium between dimeric and tetrameric forms of *Bf*GAD_WT_.

Unlike oligomeric conformations, the kinetic parameters of *Bf*GAD_WT_ vary from previously investigated prokaryotic GADs^39,44,49,51,61,62^. We find that *Bf*GAD_WT_ is an allosteric enzyme, displaying a positive cooperativity with the substrate L-glutamate. This happens when binding of substrate on one site or monomer of the protein promotes the binding of additional substrate to the other sites or monomers of the protein, where instead of a hyperbolic dependence, a sigmoidal curve is observed for the plot of initial velocity vs. substrate concentration. This phenomenon has not been seen with any other prokaryotic GAD. The physiological consequences of this cooperativity in *Bf*GAD need to be explored further. However, cooperative binding and adaptability in the allosteric enzymes helps amplify the enzyme’s response to changes in substrate concentration, making it more sensitive to the physiological substrate concentrations and conditions.

Although the kinetic parameters are different for the *Bf*GAD_WT_ compared to other studied microbial GADs, there are prominent structural similarities among all these GADs. The AlphaFold model of *Bf*GAD predicts active site architecture that is very similar to other prokaryotic GADs. Particularly, catalytic residues in the active site are positioned similar to those in *Ec*GAD and *Bt*GAD, suggesting that corresponding residues in *Bf*GAD might maintain similar interactions with ligands. While the hydrogen bonding interactions of the residues with the substrate analog glutarate are predicted to be within the expected limits of H-bond distances, in the superimposed structures of *Ec*Gad and *Bf*GAD_WT_, there are differences in the hydrogen bond lengths when amino acid substitutions are made in the active site. The changes in these H-bonding interactions could potentially be involved in some capacity in accommodating or excluding alternative substrates in the active site of the engineered *Bf*GADs to evolve new activity or specificity.

In contrast to GADs from *Bacteroides* sp., microbial GADs from various other genera have been widely characterized^39,49,61,64,65^. Most of these studies show that many annotated microbial glutamate decarboxylases prefer L-glutamate as a substrate. However, substrate promiscuity has been seen in GADs from different domains of life. For example, *E. coli* GAD was able to convert a phosphinic analog of glutamate to a phosphinic analog of GABA^58^. In addition, human and other mammalian GADs are known to catalyze decarboxylation of substrates other than L-glutamate. Specifically, sulfinic acid and sulfonic acid derivatives of alanine (known as cysteine sulfinic acid/CSA and cysteic acid/CA respectively) are known alternative substrates for mammalian GADs^57,59,66^. The decarboxylation of these molecules by mammalian GADs generate hypotaruine and taurine, from the respective sulfinic or sulfonic acids. Additionally, mammals have a couple of other *de novo* pathways for the production of taurine and hypotaurine. The primary pathway involves the enzyme cysteine sulfinic acid decarboxylase (CSAD) that generates taurine and its intermediate hypotaurine by decarboxylating either L-cysteic acid (CA) or L-cysteine sulfinic acid (CSA)^59,67^. In addition to hCSAD, mammalian GADL1 (glutamic acid decarboxylase like 1) enzyme that has very high sequence similarity with hCSAD, is capable of producing taurine^55,68^. Taurine, a sulfur containing non-proteinogenic amino acid whose production is controlled primarily by the enzyme cysteine sulfinic acid decarboxylase (CSAD)^69,70^. However, except for a few marine microbes the major *de novo* pathway for taurine formation with the help of CSAD has not been seen in other prokaryotes^57^. Interestingly, in these microbes CSAD genes are present in the operon containing cysteine dioxygenase (CDO) enzyme that catalyzes the conversion of L-cysteine to L-cysteine sulfinic acid (L-CSA) which then can be converted to hypotaurine and taurine (Supplementary Fig. 14). CDOs are not present in the members of the human gut microbiome possibly due to the hypoxic and anaerobic conditions of the gut. Despite an extensive bioinformatics search, we did not find any gut microbial genes annotated as CSAD (cysteine sulfinic acid decarboxylase). Due to the lack of the annotated CSAD enzymes responsible for the *de novo* taurine biosynthesis, we investigated the role of *Bf*GAD in the formation of taurine and its derivatives in addition to GABA.

Apart from taurine, mammalian GADL1 is also able to generate β-alanine from L-aspartate^55,68^. β-alanine is a precursor for the dipeptide beta-alanyl-L-histidine in humans, commonly known as carnosine. Carnosine is found in muscles and brain tissues at high concentrations^71,72^. In microbes, β-alanine is a precursor for coenzyme A (CoA) biosynthesis, which is an important molecule in various metabolic pathways^73^. Both carnosine and β-alanine show protective effects in individuals with cognitive deficits and Alzhiermer’s^74–76^. Moreover, higher serum concentrations of β-alanine prevent dementia^74^. Considering the structural similarities between L-glutamate and L-aspartate and the ability of some GADs to use L-aspartate as a substrate^53,56,77^, we examined the possible role of *Bf*GAD in the production of β-alanine via the decarboxylation of L-aspartate. The neuromodulatory molecules, GABA, taurine and β-alanine generated by glutamate decarboxylases or enzymes similar to GADs, have the ability to reverse cognitive deficits in neurodegenerative disorders like dementia and Alzheimer’s^12,13,24,69,74,75,78,79^. This prompted us to understand the modulation of these molecules by the members of the human gut microbiome.

Our results show that the *Bf*GAD_WT_ is promiscuous and is able to decarboxylate four additional substrates structurally similar to L-glutamate to produce hypotaurine, taurine, homotaurine, and β-alanine. There are distinct enzymes in microbes such as aspartate1-decarboxylase (A1DC) and aspartate 4-decarboxylase (A4DC) that catalyze decarboxylations of L-aspartate to produce β-alanine^80,81^ and L-alanine^82,83^ respectively. While the genome of *B. fragilis* has genes annotated for both these enzymes, it is interesting to observe that the glutamate decarboxylase from *B. fragilis* still shows decarboxylation activity towards L-aspartate to produce β-alanine. Additionally, prokaryotic A1DC and eukaryotic A1DC show evolutionary divergence where the former uses pyruvoyl cofactor whereas the latter uses PLP cofactor^84,85^. Thus, *Bf*GAD wild-type and variants harbor an activity that is mostly seen in eukaryotic organisms.

To understand the factors that drive substrate specificity in *Bf*GAD_WT_ and to evolve enzymes that potentially have switched substrate preferences or specificity, we chose two active site residues that are known to make interactions with bound ligands^39,40^. Headspace GC was used for the detection of evolved CO_2_ which is the common decarboxylation product for all enzyme catalyzed reactions (wild-type and engineered enzymes). Absolute quantification of CO_2(g)_ was challenging due to its high solubility at acidic pH where we conducted our enzymatic reactions. In these conditions, the CO_2_ produced through the reaction might still be in a soluble form as a dissolved CO_2_. So, headspace GC was utilized as a primary tool to identify alternate substrates by allowing detection of CO_2_ peak that served as a positive indication of enzymatic decarboxylation of the substrate molecules. From the identification provided by the GC experiments, subsequent analysis was carried out with TLC to detect corresponding decarboxylated products from the multiple substrates. Our headspace GC data reinforces our TLC data in most cases. Both engineered enzymes *Bf*GAD_D104N_ and *Bf*GAD_F81W_ retain preference for L-glutamate as a substrate. However, their preferences for alternate substrates are different than the *Bf*GAD_WT_. Additionally, significant spectral perturbations are observed with evolved GADs indicating that the local environment of the cofactor binding site is changed due to those amino acid substitutions. With this, we show that there might be a *de novo* pathway in gut microbes to make taurine and *Bf*GAD_WT_ is able to produce multiple neuromodulatory molecules.

## Materials and Methods

### Materials

Kanamycin, IPTG, GABase from *Pseudomonas fluorescens*, β-ME (beta-mercaptoethanol), α-ketoglutarate, D-glutamatic acid, L-cysteine sulfinic acid monohydrate, L-cysteic acid monohydrate, GABA (γ-aminobutyric acid), hypotaurine, taurine, L-aspartic acid sodium salt, β-alanine, HEPES, Imidazole, pyridoxal 5’-phosphate monohydrate and TLC Silica gel 60 F_254_ (20 cm x 20 cm) were purchased from Sigma-Aldrich. Sodium L-glutamate monohydrate, Luria-Bertani Broth (LB), buffer components, Sodium chloride, Sodium acetate, ninhydrin and NADP^+^ disodium salt were purchased from Fisher Scientific. All restriction enzymes and competent cells of *E. coli* BL21(DE3) and *E. coli* NEB5⍺ were purchased from New England Biolabs (NEB). Headspace vials were purchased from Chemglass Inc.

### Experimental procedures

#### Gene cloning

A synthesized gene for *B. fragilis* glutamate decarboxylase (*Bf*GAD) was created with the help of Genewiz from Azenta life sciences. The gene was codon optimized for expression in *E. coli* cells, which was then subcloned into a pET28a expression vector between the NdeI and HindIII restriction sites to incorporate an N-terminal hexahistidine (His_6_) Tag. The pET28a-*Bf*GAD construct was confirmed by agarose gel electrophoresis, restriction digestion, sanger sequencing (Genewiz), and full plasmid sequencing (Plasmidsaurus) and then transformed into *E. coli* NEB5⍺ and *E. coli* BL21(DE3) competent cells by heat shock. Glycerol stocks of the cells harboring the construct pET28a-*Bf*GAD were stored at –80 °C.

#### Expression and purification of *Bf*GAD_WT_ and engineered *Bf*GADs

*E. coli* BL21(DE3) cells containing the pET28a-*Bf*GADs were cultivated in LB medium containing kanamycin (50 µg/mL) with shaking (200 rpm) at 37 °C until OD_600_ reached 0.7. At this point, the *Bf*GAD expression was induced with isopropyl β-D-1-thiogalactopyranoside (IPTG) to a final concentration of 1 mM and growth was continued at 37 °C for 3 hours to allow for protein expression with continuous shaking at 200 rpm. Cells were harvested via centrifugation at 4000*g* for 30 min at 4 °C and cell pellets were stored at –80 °C. The frozen cell pellets were thawed on ice and resuspended in buffer A containing 20 mM Tris-HCl, 500 mM NaCl, 40 mM imidazole, 0.1 mM PLP, pH 7.8. All subsequent purification steps were carried out at 4 °C. The cells were disrupted by sonication (20000 Hz for 10 cycles of 30 seconds each altering 1 minute on ice) and centrifuged at 13500*g* at 4 °C for 30 min to separate supernatant from cell debris. The supernatant was then filtered through 0.2 µm PES filter membrane and loaded onto a HisTrap HP column (5 ml, 1.6 x 2.5 cm, Ni Sepharose High-performance column, GE Healthcare, now Cytiva) pre-equilibrated with buffer A at 1 mL/min. The wash step was carried out by running 5-column volumes of lysis buffer A to elute contaminating proteins. *Bf*GAD was eluted with a linear gradient from buffer A (containing 40 mM imidazole) to an elution buffer B containing 20 mM Tris-HCl, 500 mM NaCl, 400 mM imidazole, 0.1 mM PLP, pH 7.8 at 2 mL/min. Eluted protein peaks were pooled, buffer exchanged and concentrated to a final buffer (50 mM HEPES, pH 7.2) using Vivaspin 50 kDa MWCO filters (Cytiva). Aliquots of purified proteins were stored at –80 °C. Engineered *Bf*GADs were purified similarly. SDS-PAGE analysis and activity assays were performed to confirm the purity and presence of functional *Bf*GAD proteins.

#### Gel filtration chromatography with *Bf*GAD_WT_

To understand oligomeric state of *Bf*GAD_WT_, gel filtration chromatography was performed at two different pH, 4.7 (50 mM sodium acetate, 150 mM NaCl) and pH 7.2 (50 mM HEPES, 150 mM NaCl) using HiPrep™ 16/60 Sephacryl^®^ S-200 HR column. Equilibration and elution steps were carried out at a flow rate of 0.5 mL/min. High molecular weight (HMW) and Low molecular weight (LMW) calibration kits (Cytiva) were used for column calibration, to create a standard curve, and for molecular weight determination (Supplementary Fig. 5b, 5c).

### Activity assays

*Bf*GAD activity assays were performed by a coupled enzyme assay with a GABase system to measure GABA production spectrophotometrically. Briefly, *Bf*GAD was incubated with 50 mM L-glutamate in 50 mM sodium acetate, pH 4.7 buffer or in buffers at other pH values for one hour at 25 °C. Enzymatic reactions were stopped by boiling samples for 15 min. These samples were then centrifuged at 6000*g* for 5 min and if necessary, dilutions were created in 50 mM Tris-HCl, pH 8.6. For GABA measurement, 75 µL of samples (either diluted or undiluted) were added to 25 µL of Gabase assay mix containing 10 mM BME, 2 mM α-ketoglutarate, 600 µM NADP^+^, and 30 µg (0.015U/mL) of GABase in 50 mM Tris-HCl, pH 8.6^47,86–89^. In the GABase assay, GABA is converted to succinic semialdehyde (SSA) and then to succinate with subsequent production of NADPH which was measured at 340 nm. Using an extinction coefficient of 6,220 M^-1^ cm^-1^ at 340 nm, NADPH concentrations were calculated which will provide GABA concentrations in measured samples ([NADPH]=[GABA]).

### *Bf*GAD absorbance spectrophotometry and activity assays with pH variation

Spectra of *Bf*GAD_WT_ were recorded from 200 nm - 800 nm in buffers at pH 4.7, 5.5, 6.7, 7.5, and 8.6 at 25 °C on Agilent Cary 3500 UV-Vis spectrophotometer. From these spectra, absorbance changes at 335 nm (enolimine) and 420 nm (ketoenamine) for enzyme bound PLP cofactor were collected. A plot of pH versus absorbance at 420 nm was generated and the curve fitting was carried out using equation (1)^39^.

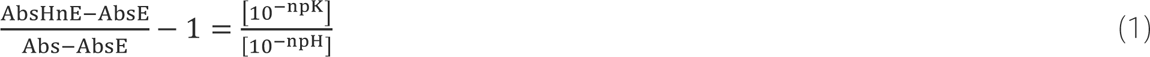

Where pK is the acid dissociation constant, number of protons involved in the titration is n and, AbsHnE and AbsE are the absorbances at acidic and basic pH which produce protonated and deprotonated forms of the enzymes. The initial activity assays for the *Bf*GAD_WT_ were performed in buffers at pH 4.7 - 8 with GABase assay system as mentioned in the above method. Based on the results of these activity assays, all subsequent enzymatic assays were carried out at pH 4.7.

### Steady-state kinetics

Kinetic parameters of *Bf*GAD_WT_ were determined by varying *Bf*GAD concentrations (0.5 µM-2 µM) and by varying L-glutamate concentrations (0.125 mM-32 mM) in 50 mM sodium acetate, pH 4.7 at 25° C. For enzyme concentration variation experiments, reaction samples with each enzyme concentration were collected at 1 min time point after starting the reaction and then stopped by boiling for 15 min. Once cooled to room temperature, these samples were analyzed for GABA content with GABase assay as mentioned above. For the substrate concentration variation experiments, 100 µL aliquots of reactions with each substrate concentration were collected at different time intervals (0.2 – 30 min). The reactions were stopped by boiling the samples for 15 minutes. Once cooled to room temperature, these samples were analyzed for GABA content with GABase assay as mentioned above. The initial velocities of various reactions were calculated by fitting data of early timepoints to a linear regression. These initial velocities were then plotted against substrate concentrations and curve fitting was carried out using equation (2).

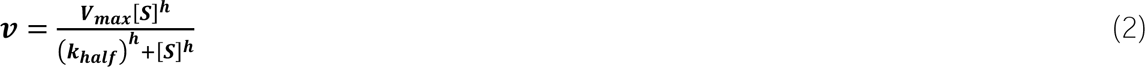

Where v is the initial velocity, V_max_ is the maximum velocity, K_half_ is the concentration of substrate at which the reaction velocity is half of V_max_, [S] is the substrate concentration, h is the hill coefficient.

### Generation of engineered *Bf*GAD variants

The pET28a-*Bf*GAD was used as a template for the generation of engineered *Bf*GAD variants. *Bf*GAD_D104N_ and *Bf*GAD_F81W_ were created by site-directed mutagenesis with Phusion DNA polymerase using the following primer pairs incorporating codons for the specific amino acid substitution (underlined).

For *Bf*GAD_D104N_

F: 5’ CCGCAATGCGCGGATATTCGGTTTCATTAATATAGTTAATG 3’

R: 5’ CATTAACTATATTAATGAAACCGAATATCCGCGCATTGCGG 3’

For *Bf*GAD_F81W_

F: 5’ CATCCATATAGGTGGTCACCCAGGTCGCCAGGTTCAGGCGCG 3’

R: 5’ CGCGCCTGAACCTGGCGACCTGGGTGACCACCTATATGGATG 3’

The variants were confirmed by gene sequencing (Genewiz) and full plasmid sequencing with Plasmidsaurus.

### CO_2(g)_ detection by headspace gas chromatography

All enzymatic reactions and controls were prepared in 6 mL vials and sealed prior to starting the reaction. The reactions were started by the addition of *Bf*GAD (WT or engineered variants) with a syringe into the reaction mixture containing 50 mM substrate in 50 mM sodium acetate, pH 4.7. These vials were incubated at 37 °C for 24 hours and 48 hours. Headspace GC measurements were carried out on an Agilent 8890 gas chromatograph system equipped with a flame ionization detector (FID) and a Hayesep Q packed column (1.8 m x 2 mm x 3.17 mm) (Agilent) operating with an Argon carrier gas (flow rate= 5 mL/min). The oven was programed to hold 30 °C for 6.5 min, ramp at 30 °C/min to 280 °C with a hold for 4 min for a total run time of 18.83 min. The flame ionization detector was used for the detection of CO_2_ gas with a temperature setting of 275 °C, hydrogen flow of 60 mL/min, air flow of 400 mL/min and constant makeup gas (nitrogen) at 5 mL/min. The retention time for CO_2_ gas was 12.83 min. Multiple standards of CO_2_ gas were analyzed by this method to confirm retention time before injecting reaction samples (Supplementary Fig. 15).

### Thin layer chromatography (TLC)

Silica gel plates (stationary phase) with a solvent system (mobile phase) of 3:1:1 ratio of butanol: acetic acid: H_2_O were used to separate reaction products from substrates^60^. 2 µL of reaction mixtures were spotted on glass silica plates along with 10 mM of various metabolites (L-glutamate, D-glutamate, L-CSA, L-CA, L-HCA, L-Aspartate, GABA, hypotaurine, taurine, homotaurine and β-alanine) as standards. Separation via TLC was carried out at 25 °C for 3-4 hours in an above-mentioned mobile phase. Once mobile phase reached a sufficient height on the TLC plate, chromatographic separation was discontinued. Plates were treated with 0.5% ninhydrin in acetone (w/v) and heated minimally with a dryer for the color development. The TLC plate images were initially inverted via ImageJ and then intensity of each spot was quantified by a custom python code.

### Liquid chromatography with tandem mass spectrometry (LC-MS/MS) detection of decarboxylated products from reactions catalyzed by *Bf*GADs (WT and engineered variants)

Samples were analyzed by ESI-LC-MS/MS in positive (products) and negative ion (substrates) mode using a Thermo vanquish LC and TSQ Altis Triple Quadrupole Mass Spectrometer. Samples were separated by gradient elution using Agilent Infinity lab Poroshell 120 HILIC-Z 2.1 x 100 mm, 2.7 µm column at 25 °C with 20 mM ammonium formate, pH 3.0 (Solvent A) and 20 mM ammonium formate in 9:1 acetonitrile: H_2_O, pH 3.0 (Solvent B). Elution was initiated at 70% B for 11.5 min, followed by gradient elution from 70 to 100% B over 11.5 min, and 100% B for 4 min at a flow rate of 0.5mL/min. The precursor to product transitions of *m/z* [M+H]^-^152→ 88.1(Cysteine sulfinic acid), [M+H]^-^ 168→ 81.1(Cysteic acid), [M+H]^+^110.1→ 45.1(hypotaurine), [M+H]^+^126.1→ 44.1(taurine) were employed to monitor substrates and products.

### Multiple Sequence alignment and phylogenetic analyses of gut microbial GADs

All the protein sequences were obtained from NCBI. Multiple sequence alignment was performed using Clustal Omega (1.2.4) tool with default parameters^90,91^ to understand sequence similarities among various gut microbial glutamate decarboxylases (GADs). The phylogenetic tree was created for *Bacteroides* GADs using cyberinfrastructure for phylogenetic research (CIPRES)^92^. Within CIPRES, the MAFFT on XSEDE (7.505)^93^ was utilized to create a separate multiple sequence alignment that goes through FastTreeMP on XSEDE(2.1.10)^94,95^ to obtain the phylogenetic tree file which can be visualized and annotated with Interactive tree of life (iTOL v6.8.1)^96^.

### *Bf*GAD dimer generation with the AlphaFold2 and visualization, structural alignments, and mutagenesis via PyMOL

The dimer structure of wildtype *Bf*GAD was generated using AlphaFold2 and visualized using PyMOL. Variants were then created using the mutagenesis wizard in PyMOL. Rotamers for the altered residues were selected based on their best fit to the position of wild-type residues in the *Bf*GAD protein. All structural alignments were carried out using PyMOL align method, involving five iteration cycles and a cutoff of 2 Å.

## Acknowledgments

This research is supported by the seed grant from Edson initiative for the dementia care and solutions to DDS. We acknowledge resources and support from the Knowledge Enterprise Biosciences Core Facilities at Arizona State University. We thank Dr. Rosa Krajmalnik-Brown and Christopher Muse from the Biodesign Centre for Health Through Microbiomes for the help with headspace GC. This research is supported by the seed grant from Edson initiative for the dementia care and solutions to DDS. AS was supported by an NIH-NIGMS MIRA award R35GM147131.

## Author Contributions

PD performed majority of the experiments and data analysis. CWP performed some of the TLC experiments. AS performed analysis and data fitting for pH and Abs variation experiment and provided code and advice on the image analysis of TLC plates, PD and DDS conceptualized the project and wrote the paper.

## Data availability

All the data are available in the manuscript or with associated supporting information. The python code used for TLC image analysis is available at https://github.com/Pavani-dadi/TLC_profile.

---

**Scheme 1.**
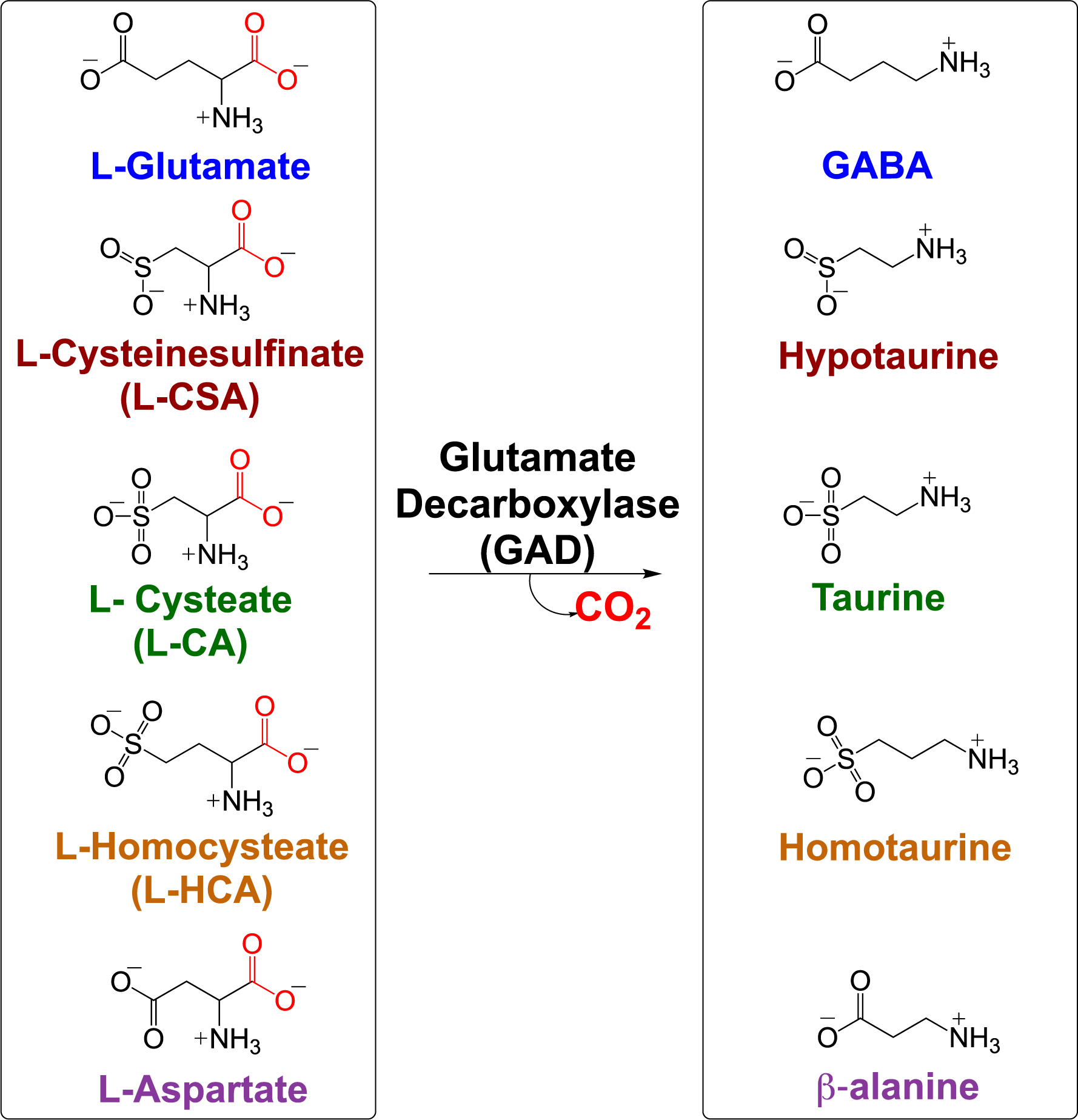
Predicted substrates and products of *Bf*GAD catalyzed decarboxylations.

## References

1 Murley, A. G. et al. GABA and glutamate deficits from frontotemporal lobar degeneration are associated with disinhibition. Brain 143, 3449–3462 (2020). 10.1093/brain/awaa305

2 Nuss, P. Anxiety disorders and GABA neurotransmission: a disturbance of modulation. Neuropsychiatr Dis Treat 11, 165–175 (2015). 10.2147/NDT.S58841

3 Cawley, N. et al. Reduced gamma-aminobutyric acid concentration is associated with physical disability in progressive multiple sclerosis. Brain 138, 2584–2595 (2015). 10.1093/brain/awv209

4 Goddard, A. W. et al. Reductions in occipital cortex GABA levels in panic disorder detected with 1h-magnetic resonance spectroscopy. Arch Gen Psychiatry 58, 556–561 (2001). 10.1001/archpsyc.58.6.556

5 Goddard, A. W. et al. Impaired GABA neuronal response to acute benzodiazepine administration in panic disorder. Am J Psychiatry 161, 2186–2193 (2004). 10.1176/appi.ajp.161.12.2186

6 Long, Z. et al. Decreased GABA levels in anterior cingulate cortex/medial prefrontal cortex in panic disorder. Prog Neuropsychopharmacol Biol Psychiatry 44, 131–135 (2013). 10.1016/j.pnpbp.2013.01.020

7 Orhan, F. et al. CSF GABA is reduced in first-episode psychosis and associates to symptom severity. Mol Psychiatry 23, 1244–1250 (2018). 10.1038/mp.2017.25

8 Sarlo, G. L. & Holton, K. F. Brain concentrations of glutamate and GABA in human epilepsy: A review. Seizure 91, 213–227 (2021). 10.1016/j.seizure.2021.06.028

9 Umesawa, Y. et al. Altered GABA Concentration in Brain Motor Area Is Associated with the Severity of Motor Disabilities in Individuals with Autism Spectrum Disorder. J Autism Dev Disord 50, 2710–2722 (2020). 10.1007/s10803-020-04382-x

10 Porges, E. C. et al. Frontal Gamma-Aminobutyric Acid Concentrations Are Associated With Cognitive Performance in Older Adults. Biol Psychiatry Cogn Neurosci Neuroimaging 2, 38–44 (2017). 10.1016/j.bpsc.2016.06.004

11 Carello-Collar, G. et al. The GABAergic system in Alzheimer’s disease: a systematic review with meta-analysis. Mol Psychiatry (2023). 10.1038/s41380-023-02140-w

12 Solas, M., Puerta, E. & Ramirez, M. Treatment Options in Alzheimeŕs Disease: The GABA Story. Current Pharmaceutical Design 21, 4960–4971 (2015). 10.2174/1381612821666150914121149

13 Jia, F. et al. Taurine is a potent activator of extrasynaptic GABA(A) receptors in the thalamus. J Neurosci 28, 106–115 (2008). 10.1523/JNEUROSCI.3996-07.2008

14 Ochoa-de la Paz, L., Zenteno, E., Gulias-Cañizo, R. & Quiroz-Mercado, H. Taurine and GABA neurotransmitter receptors, a relationship with therapeutic potential? Expert Review of Neurotherapeutics 19, 289–291 (2019). 10.1080/14737175.2019.1593827

15 Alom, J., Mahy, J. N., Brandi, N. & Tolosa, E. Cerebrospinal fluid taurine in Alzheimer’s disease. Ann Neurol 30, 735 (1991). 10.1002/ana.410300518

16 Aytan, N. et al. Fingolimod modulates multiple neuroinflammatory markers in a mouse model of Alzheimer’s disease. Sci Rep 6, 24939 (2016). 10.1038/srep24939

17 Chiquita, S. et al. A longitudinal multimodal in vivo molecular imaging study of the 3xTg-AD mouse model shows progressive early hippocampal and taurine loss. Hum Mol Genet 28, 2174–2188 (2019). 10.1093/hmg/ddz045

18 Garcia-Serrano, A. M., Vieira, J. P. P., Fleischhart, V. & Duarte, J. M. N. Taurine and N-acetylcysteine treatments prevent memory impairment and metabolite profile alterations in the hippocampus of high-fat diet-fed female mice. Nutr Neurosci 26, 1090–1102 (2023). 10.1080/1028415X.2022.2131062

19 Pomara, N. et al. Glutamate and other CSF amino acids in Alzheimer’s disease. Am J Psychiatry 149, 251–254 (1992). 10.1176/ajp.149.2.251

20 Vermeiren, Y. et al. Behavioral correlates of cerebrospinal fluid amino acid and biogenic amine neurotransmitter alterations in dementia. Alzheimers Dement 9, 488–498 (2013). 10.1016/j.jalz.2012.06.010

21 Csernansky, J. G., Bardgett, M. E., Sheline, Y. I., Morris, J. C. & Olney, J. W. CSF excitatory amino acids and severity of illness in Alzheimer’s disease. Neurology 46, 1715–1720 (1996). 10.1212/wnl.46.6.1715

22 Gao, R., Bae, M. A., Chang, K. J. & Kim, S. H. Comparison of Urinary Excretion of Taurine Between Elderly with Dementia and Normal Elderly. Adv Exp Med Biol 975 Pt 1, 57–65 (2017). 10.1007/978-94-024-1079-2_5

23 Fujii, Y. et al. Fecal metabolite of a gnotobiotic mouse transplanted with gut microbiota from a patient with Alzheimer’s disease. Biosci Biotechnol Biochem 83, 2144–2152 (2019). 10.1080/09168451.2019.1644149

24 Jang, H. et al. Taurine Directly Binds to Oligomeric Amyloid-beta and Recovers Cognitive Deficits in Alzheimer Model Mice. Adv Exp Med Biol 975 Pt 1, 233–241 (2017). 10.1007/978-94-024-1079-2_21

25 Kim, H. Y. et al. Taurine in drinking water recovers learning and memory in the adult APP/PS1 mouse model of Alzheimer’s disease. Sci Rep 4, 7467 (2014). 10.1038/srep07467

26 Manzano, S., Aguera, L., Aguilar, M. & Olazaran, J. A Review on Tramiprosate (Homotaurine) in Alzheimer’s Disease and Other Neurocognitive Disorders. Front Neurol 11, 614 (2020). 10.3389/fneur.2020.00614

27 Singh, P. et al. Taurine deficiency as a driver of aging. Science 380, eabn9257 (2023). 10.1126/science.abn9257

28 Borsom, E. M., Lee, K. & Cope, E. K. Do the Bugs in Your Gut Eat Your Memories? Relationship between Gut Microbiota and Alzheimer’s Disease. Brain Sci 10 (2020). 10.3390/brainsci10110814

29 Saji, N. et al. Analysis of the relationship between the gut microbiome and dementia: a cross-sectional study conducted in Japan. Sci Rep 9, 1008 (2019). 10.1038/s41598-018-38218-7

30 Zhuang, Z.-Q. et al. Gut Microbiota is Altered in Patients with Alzheimer’s Disease. Journal of Alzheimer’s Disease 63, 1337–1346 (2018). 10.3233/JAD-180176

31 Guo, M. et al. Gut Microbiome Features of Chinese Patients Newly Diagnosed with Alzheimer’s Disease or Mild Cognitive Impairment. Journal of Alzheimer’s Disease 80, 299–310 (2021). 10.3233/JAD-201040

32 Li, B. et al. Mild cognitive impairment has similar alterations as Alzheimer’s disease in gut microbiota. Alzheimer’s & Dementia 15, 1357–1366 (2019). 10.1016/j.jalz.2019.07.002

33 Jemimah, S., Chabib, C. M. M., Hadjileontiadis, L. & AlShehhi, A. Gut microbiome dysbiosis in Alzheimer’s disease and mild cognitive impairment: A systematic review and meta-analysis. PLOS ONE 18, e0285346 (2023). 10.1371/journal.pone.0285346

34 Salyers, A. A. Bacteroides of the human lower intestinal tract. Annu Rev Microbiol 38, 293–313 (1984). 10.1146/annurev.mi.38.100184.001453

35 Wexler, H. M. Bacteroides: the good, the bad, and the nitty-gritty. Clin Microbiol Rev 20, 593–621 (2007). 10.1128/CMR.00008-07

36 Strandwitz, P. et al. GABA-modulating bacteria of the human gut microbiota. Nat Microbiol 4, 396–403 (2019). 10.1038/s41564-018-0307-3

37 Otaru, N. et al. GABA Production by Human Intestinal Bacteroides spp.: Prevalence, Regulation, and Role in Acid Stress Tolerance. Frontiers in Microbiology 12, 656895 (2021). 10.3389/fmicb.2021.656895

38 Horvath, T. D. et al. Bacteroides ovatus colonization influences the abundance of intestinal short chain fatty acids and neurotransmitters. iScience 25, 104158 (2022). 10.1016/j.isci.2022.104158

39. Capitani, G. et al. Crystal structure and functional analysis of *Escherichia coli* glutamate decarboxylase. The EMBO Journal 22, 4027-4037-4037 (2003). 10.1093/emboj/cdg403

40 Dutyshev, D. I. et al. Structure of *Escherichia coli* glutamate decarboxylase (GADα) in complex with glutarate at 2.05 Å resolution. Acta Crystallographica Section D Biological Crystallography 61, 230-235 (2005). 10.1107/S0907444904032147

41 Beattie, A. E. et al. The Pyridoxal 5′-Phosphate (PLP)-Dependent Enzyme Serine Palmitoyltransferase (SPT): Effects of the Small Subunits and Insights from Bacterial Mimics of Human hLCB2a HSAN1 Mutations. BioMed Research International 2013, 1–13 (2013). 10.1155/2013/194371

42 Moore, P. S., Dominici, P. & Borri Voltattorni, C. Cloning and expression of pig kidney dopa decarboxylase: comparison of the naturally occurring and recombinant enzymes. Biochemical Journal 315, 249–256 (1996). 10.1042/bj3150249

43 Zhou, X. & Toney, M. D. pH Studies on the Mechanism of the Pyridoxal Phosphate-Dependent Dialkylglycine Decarboxylase. Biochemistry 38, 311–320 (1999). 10.1021/bi981455s

44 Liu, S. et al. Coordinated regulation of Bacteroides thetaiotaomicron glutamate decarboxylase activity by multiple elements under different pH. Food Chem 403, 134436 (2023). 10.1016/j.foodchem.2022.134436

45 Pennacchietti, E. et al. Mutation of His465 Alters the pH-dependent Spectroscopic Properties of Escherichia coli Glutamate Decarboxylase and Broadens the Range of Its Activity toward More Alkaline pH. Journal of Biological Chemistry 284, 31587–31596 (2009). 10.1074/jbc.M109.049577

46 Shukuya, R. & Schwert, G. W. Glutamic acid decarboxylase. II. The spectrum of the enzyme. J Biol Chem 235, 1653–1657 (1960).

47 Tramonti, A., John, R. A., Bossa, F. & De Biase, D. Contribution of Lys276 to the conformational flexibility of the active site of glutamate decarboxylase from *Escherichia coli*: Role of Lys276 in E. coli glutamate decarboxylase. European Journal of Biochemistry 269, 4913-4920 (2002). 10.1046/j.1432-1033.2002.03149.x

48 De Biase, D. & Pennacchietti, E. Glutamate decarboxylase-dependent acid resistance in orally acquired bacteria: function, distribution and biomedical implications of the gadBC operon. Mol Microbiol 86, 770–786 (2012). 10.1111/mmi.12020

49 Chang, C. et al. Purification and characterization of glutamate decarboxylase from Enterococcus raffinosus TCCC11660. J Ind Microbiol Biotechnol 44, 817–824 (2017). 10.1007/s10295-017-1906-3

50 Fonda, M. L. Glutamate decarboxylase. Substrate specificity and inhibition by carboxylic acids. Biochemistry 11, 1304–1309 (1972). 10.1021/bi00757a029

51 Ueno, Y., Hayakawa, K., Takahashi, S. & Oda, K. Purification and characterization of glutamate decarboxylase from Lactobacillus brevis IFO 12005. Biosci Biotechnol Biochem 61, 1168–1171 (1997). 10.1271/bbb.61.1168

52 Fonda, M. L. L-Glutamate decarboxylase from bacteria. Methods Enzymol 113, 11–16 (1985). 10.1016/s0076-6879(85)13005-3

53 Kim, H. W., Kashima, Y., Ishikawa, K. & Yamano, N. Purification and characterization of the first archaeal glutamate decarboxylase from Pyrococcus horikoshii. Biosci Biotechnol Biochem 73, 224–227 (2009). 10.1271/bbb.80583

54 Tomita, H., Yokooji, Y., Ishibashi, T., Imanaka, T. & Atomi, H. An archaeal glutamate decarboxylase homolog functions as an aspartate decarboxylase and is involved in beta-alanine and coenzyme A biosynthesis. J Bacteriol 196, 1222–1230 (2014). 10.1128/JB.01327-13

55 Liu, P. et al. Role of glutamate decarboxylase-like protein 1 (GADL1) in taurine biosynthesis. J Biol Chem 287, 40898–40906 (2012). 10.1074/jbc.M112.393728

56 Richardson, G. et al. An examination of aspartate decarboxylase and glutamate decarboxylase activity in mosquitoes. Mol Biol Rep 37, 3199–3205 (2010). 10.1007/s11033-009-9902-y

57 Agnello, G., Chang, L. L., Lamb, C. M., Georgiou, G. & Stone, E. M. Discovery of a Substrate Selectivity Motif in Amino Acid Decarboxylases Unveils a Taurine Biosynthesis Pathway in Prokaryotes. ACS Chemical Biology 8, 2264–2271 (2013). 10.1021/cb400335k

58 Daniela De Biase, F. C., Eugenia Pennacchietti, Fabio Giovannercole, Antonio Coluccia, Jouko Vepsäläinen & Alex Khomutov Enzymatic kinetic resolution of desmethylphosphinothricin indicates that phosphinic group is a bioisostere of carboxyl group. Commnications chemistry (2020).

59 Wu, J. Y. Purification and characterization of cysteic acid and cysteine sulfinic acid decarboxylase and L-glutamate decarboxylase from bovine brain. Proc Natl Acad Sci U S A 79, 4270–4274 (1982). 10.1073/pnas.79.14.4270

60 Tramonti, A. et al. A Novel, Easy Assay Method for Human Cysteine Sulfinic Acid Decarboxylase. Life 11, 438 (2021). 10.3390/life11050438

61 Komatsuzaki, N., Nakamura, T., Kimura, T. & Shima, J. Characterization of glutamate decarboxylase from a high gamma-aminobutyric acid (GABA)-producer, Lactobacillus paracasei. Biosci Biotechnol Biochem 72, 278–285 (2008). 10.1271/bbb.70163

62 Hiraga, K., Ueno, Y. & Oda, K. Glutamate decarboxylase from Lactobacillus brevis: activation by ammonium sulfate. Biosci Biotechnol Biochem 72, 1299–1306 (2008). 10.1271/bbb.70782

63 Giovannercole, F. et al. On the effect of alkaline pH and cofactor availability in the conformational and oligomeric state of Escherichia coli glutamate decarboxylase. Protein Eng Des Sel 30, 235–244 (2017). 10.1093/protein/gzw076

64 Lee, J. Y. & Jeon, S. J. Characterization and immobilization on nickel-chelated Sepharose of a glutamate decarboxylase A from Lactobacillus brevis BH2 and its application for production of GABA. Biosci Biotechnol Biochem 78, 1656–1661 (2014). 10.1080/09168451.2014.936347

65 Nomura, M. et al. Lactococcus lactis contains only one glutamate decarboxylase gene. Microbiology (Reading) 145 (Pt 6), 1375–1380 (1999). 10.1099/13500872-145-6-1375

66 Blindermann, J. M., Maitre, M., Ossola, L. & Mandel, P. Purification and some properties of L-glutamate decarboxylase from human brain. Eur J Biochem 86, 143–152 (1978). 10.1111/j.1432-1033.1978.tb12293.x

67 Winge, I. et al. Mammalian CSAD and GADL1 have distinct biochemical properties and patterns of brain expression. Neurochemistry International 90, 173–184 (2015). 10.1016/j.neuint.2015.08.013

68 Mahootchi, E. et al. GADL1 is a multifunctional decarboxylase with tissue-specific roles in beta-alanine and carnosine production. Sci Adv 6, eabb3713 (2020). 10.1126/sciadv.abb3713

69 Ripps, H. & Shen, W. Review: taurine: a “very essential“ amino acid. Molecular Vision 18, 2673–2686 (2012).

70 De La Rosa, J. & Stipanuk, M. H. Evidence for a rate-limiting role of cysteinesulfinate decarboxylase activity in taurine biosynthesis in vivo. Comparative Biochemistry and Physiology Part B: Comparative Biochemistry 81, 565–571 (1985). 10.1016/0305-0491(85)90367-0

71 Boldyrev, A. A., Aldini, G. & Derave, W. Physiology and pathophysiology of carnosine. Physiol Rev 93, 1803–1845 (2013). 10.1152/physrev.00039.2012

72 Sale, C., Saunders, B. & Harris, R. C. Effect of beta-alanine supplementation on muscle carnosine concentrations and exercise performance. Amino Acids 39, 321–333 (2010). 10.1007/s00726-009-0443-4

73 Lopez-Samano, M. et al. A novel way to synthesize pantothenate in bacteria involves beta-alanine synthase present in uracil degradation pathway. Microbiologyopen 9, e1006 (2020). 10.1002/mbo3.1006

74 Hata, J. et al. Association Between Serum beta-Alanine and Risk of Dementia. Am J Epidemiol 188, 1637–1645 (2019). 10.1093/aje/kwz116

75 Ostfeld, I. et al. Role of &beta;-Alanine Supplementation on Cognitive Function, Mood, and Physical Function in Older Adults; Double-Blind Randomized Controlled Study. Nutrients 15 (2023).

76 Dhingra, D., Parle, M. & Kulkarni, S. K. β - Alanine protects mice from memory deficits induced by ageing, scopolamine, diazepam and ethanol. Indian Journal of Pharmaceutical Sciences 68 (2006). 10.4103/0250-474X.25718

77 Wu, J.-Y., Matsuda, T. & Roberts, E. Purification and Characterization of Glutamate Decarboxylase from Mouse Brain. Journal of Biological Chemistry 248, 3029–3034 (1973). 10.1016/S0021-9258(19)44004-0

78 Albrecht, J. & Schousboe, A. Taurine Interaction with Neurotransmitter Receptors in the CNS: An Update. Neurochemical Research 30, 1615–1621 (2005). 10.1007/s11064-005-8986-6

79 Bae, M. A., Lee, E. S., Cho, S. M., Kim, S. H. & Chang, K. J. The Effects of Dietary Taurine-Containing Jelly Supplementation on Cognitive Function and Memory Ability of the Elderly with Subjective Cognitive Decline. Adv Exp Med Biol 1370, 395–403 (2022). 10.1007/978-3-030-93337-1_37

80 Cui, W., Liu, H., Ye, Y., Han, L. & Zhou, Z. Discovery and Engineering of a Novel Bacterial L-Aspartate alpha-Decarboxylase for Efficient Bioconversion. Foods 12 (2023). 10.3390/foods12244423

81 Begley, T. P., Kinsland, C. & Strauss, E. The biosynthesis of coenzyme A in bacteria. Vitam Horm 61, 157–171 (2001). 10.1016/s0083-6729(01)61005-7

82 Lima, S. et al. The crystal structure of the Pseudomonas dacunhae aspartate-beta-decarboxylase dodecamer reveals an unknown oligomeric assembly for a pyridoxal-5’-phosphate-dependent enzyme. J Mol Biol 388, 98–108 (2009). 10.1016/j.jmb.2009.02.055

83 Jha, S., Speth, R. C. & Macheroux, P. The possible role of a bacterial aspartate beta-decarboxylase in the biosynthesis of alamandine. Med Hypotheses 144, 110038 (2020). 10.1016/j.mehy.2020.110038

84 Yu, X. J. et al. Protein Engineering of a Pyridoxal-5’-Phosphate-Dependent l-Aspartate-alpha-Decarboxylase from Tribolium castaneum for beta-Alanine Production. Molecules 25 (2020). 10.3390/molecules25061280

85 Webb, M. E. et al. Structure of Escherichia coli aspartate alpha-decarboxylase Asn72Ala: probing the role of Asn72 in pyruvoyl cofactor formation. Acta Crystallogr Sect F Struct Biol Cryst Commun 68, 414–417 (2012). 10.1107/S1744309112009487

86 Cozzani, I. Spectrophotometric assay of l-glutamic acid decarboxylase. Analytical Biochemistry 33, 125–131 (1970). 10.1016/0003-2697(70)90446-X

87 Tramonti, A., De Biase, D., Giartosio, A., Bossa, F. & John, R. A. The roles of His-167 and His-275 in the reaction catalyzed by glutamate decarboxylase from Escherichia coli. J Biol Chem 273, 1939–1945 (1998). 10.1074/jbc.273.4.1939

88 Tsukatani, T., Higuchi, T. & Matsumoto, K. Enzyme-based microtiter plate assay for γ-aminobutyric acid: Application to the screening of γ-aminobutyric acid-producing lactic acid bacteria. Analytica Chimica Acta 540, 293–297 (2005). 10.1016/j.aca.2005.03.056

89 De Biase, D., Tramonti, A., John, R. A. & Bossa, F. Isolation, overexpression, and biochemical characterization of the two isoforms of glutamic acid decarboxylase from Escherichia coli. Protein Expr Purif 8, 430–438 (1996). 10.1006/prep.1996.0121

90 Goujon, M. et al. A new bioinformatics analysis tools framework at EMBL-EBI. Nucleic Acids Res 38, W695–699 (2010). 10.1093/nar/gkq313

91 Sievers, F. et al. Fast, scalable generation of high-quality protein multiple sequence alignments using Clustal Omega. Mol Syst Biol 7, 539 (2011). 10.1038/msb.2011.75

92 Miller, M. A., Pfeiffer, W. & Schwartz, T. in 2010 Gateway Computing Environments Workshop (GCE). 1–8.

93 Katoh, K., Misawa, K., Kuma, K. & Miyata, T. MAFFT: a novel method for rapid multiple sequence alignment based on fast Fourier transform. Nucleic Acids Res 30, 3059–3066 (2002). 10.1093/nar/gkf436

94 Price, M. N., Dehal, P. S. & Arkin, A. P. FastTree: computing large minimum evolution trees with profiles instead of a distance matrix. Mol Biol Evol 26, 1641–1650 (2009). 10.1093/molbev/msp077

95 Price, M. N., Dehal, P. S. & Arkin, A. P. FastTree 2--approximately maximum-likelihood trees for large alignments. PLoS One 5, e9490 (2010). 10.1371/journal.pone.0009490

96 Letunic, I. & Bork, P. Interactive Tree Of Life (iTOL) v5: an online tool for phylogenetic tree display and annotation. Nucleic Acids Res 49, W293–W296 (2021). 10.1093/nar/gkab301

